# Pre-anthesis spike growth dynamics and its association to yield components among elite bread wheat cultivars (*Triticum aestivum* L. spp.) under Mediterranean climate

**DOI:** 10.1101/2023.01.08.523132

**Authors:** Rajib Roychowdhury, Orian Zilberman, Kottakota Chandrasekhar, Arie Curzon, Kamal Nashef, Shahal Abbo, Gustavo A. Slafer, David J. Bonfil, Roi Ben-David

**Affiliations:** Department of Vegetables and Field Crops, The Institute of Plant Sciences, Agriculture Research Organization (ARO) – Volcani Center, Rishon Lezion, Israel; The Robert H. Smith Institute of Plant Science and Genetics in Agriculture, The Levi Eshkol School of Agriculture, The Hebrew University of Jerusalem, Rehovot, Israel; Department of Entomology, Institute of Plant Protection, Agriculture Research Organization (ARO) – Volcani Center, Rishon Lezion, Israel; Department of Crop and Forest Sciences, University of Lleida - AGROTECNIO Center, Av. Rovira Roure 191, 25198 Lleida, Spain; Agricultural Research Organization (ARO), Gilat Research Center, M.P. Negev, Israel

**Keywords:** Pre-anthesis, Terminal spikelet, Stem elongation, Yield components, *Triticum aestivum*

## Abstract

Wheat (*Triticum* spp.) grain yield (GY) is highly associated with grain number per unit area (GN m^-2^). Biomass accumulation and partitioning are essential to understand pre-anthesis spike growth dynamics which determines spike dry matter at anthesis (SDMa) - a GN determinant. Spike growth takes place during the stem elongation period (SE), from terminal spikelet to anthesis, following leaf and spikelet initiation (LS) from sowing to terminal spikelet. In this study, bread wheat cultivars were examined under Mediterranean semi-arid conditions to determine (i) the varietal differences in pre-anthesis phase duration, (ii) whether this variability influences biomass partitioning and spike-related traits, and (iii) to what extent, the genotypic variations in pre-anthesis phase duration and spike growth are associated with yield components. A panel of Israeli commercial bread wheat cultivars were grown in the field during 2016-17 (three environments) and 2017-18 (two environments) and characterized for pre-anthesis phases, floral conditions and spike fertility *via* histological measurements, spike traits and dry matter accumulation and partitioning at anthesis and maturity and for yield components. Significant variability in the timing of pre-anthesis phases was detected within the tested panel. LS duration, and occasionally SE, favourably related with a better dry matter of fertile florets spike^-1^ (at anthesis) and SDM (at both anthesis and maturity). Two cultivar pairs ‘Zahir-Yuval’ and ‘Negev-Gedera’, which flowered concurrently, revealed significant differences in the durations of LS and SE phases across the environments. Longer LS (e.g., in cultivars Zahir and Negev) exhibited increased spikelets number spike^-1^, whereas longer SE (e.g., in Yuval and Gedera) enhanced spike fertility through improving the survival rate of floret primordia (FSR%) of central spikelets. However, there was a trade-off for FSR at the proximal and distal spike portions, resulting reduction of final GN (or GY) in cultivars with longer SE. It is concluded that, in this panel, the duration of both LS and SE contribute to spike fertility. However, under short wheat growing cycle, LS duration seemed a stronger driver than SE for GN and yield enhancement. These highlights the novel importance of pre-anthesis phases, especially the role of LS in wheat yield increment during the short growing cycle. The varietal combination with variable LS and SE duration could be implemented in the breeding pipeline and used as pre-breeding materials for GN improvement. Furthermore, the findings will improve pre-anthesis traits adoption in Mediterranean bread wheat future breeding programs.

## 1. Introduction

Wheat (*Triticum* spp.) is one of the major globally important food crops, in terms of cultivated area and food supply, with a worldwide annual production of 760 million tons across 219 million hectares (FAOSTAT, 2021). Bread wheat (*Triticum aestivum* L) presently dominates about 65% of total wheat growing areas due to its increased production, adaptability in a wide range of geographical areas, and marketability (Reynolds et al., 2012). The dryland semi-arid Mediterranean climate of Israel is characterized by winter rainfall, a relatively short wheat-growing season (from mid-November to May) with increasing temperatures and a lack of sufficient soil moisture at the grain-filling stage (Amram et al., 2015). The terminal drought and spring heat waves have negative effects on grain yield (GY) and quality (Telfer et al., 2018) and this generally dictates clear advantage to early flowering (anthesis) cultivars (Loss and Siddique, 1994). However, the situation may be more complicated in the Mediterranean regions in general; although still true that late flowering must be avoided, it may not necessarily be true all the time (Savin et al., 2015). In general, the anthesis time of wheat is controlled by genetic and environmental factors like photoperiod (day-length) and vernalization (accumulating low-temperature to promote anthesis) to adapt the crop in various geographical locations (Gomez et al., 2014). Israeli wheat cultivars are mostly insensitive to both photoperiod and vernalization and therefore the differences between them in time to anthesis must be due to *earliness per se* (*eps*), and indeed *eps* may reflect differences in sensitivity to temperature (Slafer et al., 2021).

Yield is mostly sink-limited during grain-filling, i.e., the available photosynthates, including post-anthesis actual photosynthesis plus reserves stored pre-anthesis as water-soluble carbohydrates in vegetative tissues, are more than the demand of developing grains in the spike (González et al., 2014; Reynolds et al., 2022; Serrago et al., 2013). Therefore, wheat GY highly depends far more on the number of grains per m^2^ (GN m^-2^) than on their average weight (GW) (Sadras and Slafer, 2012; Slafer et al., 2014). Final GN m^-2^ is a combination of yield components such as the number of plants m^-2^, number of fertile tillers per unit area, number of spikelets spike^-1^ and grains spikelet^-1^, which depends largely on the floret survival rate (FSR%) (Slafer et al., 2009). As the processes determining each of these components of GN m^-2^ take place during the stem elongation (SE) phase of pre-anthesis development (Slafer et al., 2021), an increase in dry matter (DM) partitioning to developing spikes during pre-anthesis might contribute to optimizing the GY (Elía et al., 2016; Serrago et al., 2013; Terrile et al., 2017). As late floret development (i.e., the survival of floret primordia that determines the number of fertile florets; e.g. Ferrante et al., 2020) is limited by the source strength (Ferrante et al., 2010; Reynolds et al., 2022), increasing spike growth before anthesis would result in improvements of spike fertility. This is the physiological basis for the widely reported positive relationship between GN m^-2^ and spike dry matter at anthesis (SDMa) (Abbate et al., 2013; Basavaraddi et al., 2021; Calderini et al., 1999; Ephrat, 1974; Slafer and Andrade, 1993). Slafer and Andrade (1993) suggested that increased GN m^-2^ of bread wheat may be due to not only increased allocation of assimilates to the juvenile spikes before anthesis, but also the partitioning within the spike assigning a higher fraction of SDM to the developing florets rather than the structural parts of the spike, like rachis, glume, lemma and paleas, being this a likely source of improvements in fruiting efficiency (Slafer et al., 2015).

The semi-dwarf varieties were characterized by prolific spikes as a result of the reduced competition of assimilates within the plant (Ephrat, 1974). In order to further reduce the competition of assimilates between the developing spike and growing stem, fine-tuning of pre-anthesis developmental stages, especially those occurring before and after the onset of SE, was suggested as a potential means for GN improvement (Beche et al., 2018; Slafer, 2012; Slafer and Rawson, 1994). The developmental cycle of pre-anthesis spike growth comprises two major phases: leaf and spikelet initiation phase (LS) ranging from sowing (SW) to terminal spikelet (TS), and that of stem elongation (SE), from TS to anthesis (AN). LS starts at sowing (SW) and ends at terminal spikelet (TS), i.e. the differentiation of the shoot apical meristem, which includes the vegetative phase - leaf initiation (from sowing to floral initiation; FI) and early reproductive phase of spikelet initiation (from FI to TS stage). The late-reproductive SE phase starts at TS (or actual onset of stem elongation of the main stem as a phenotypic marker) and ends at anthesis (for details see Figure 1).

**Figure 1.**
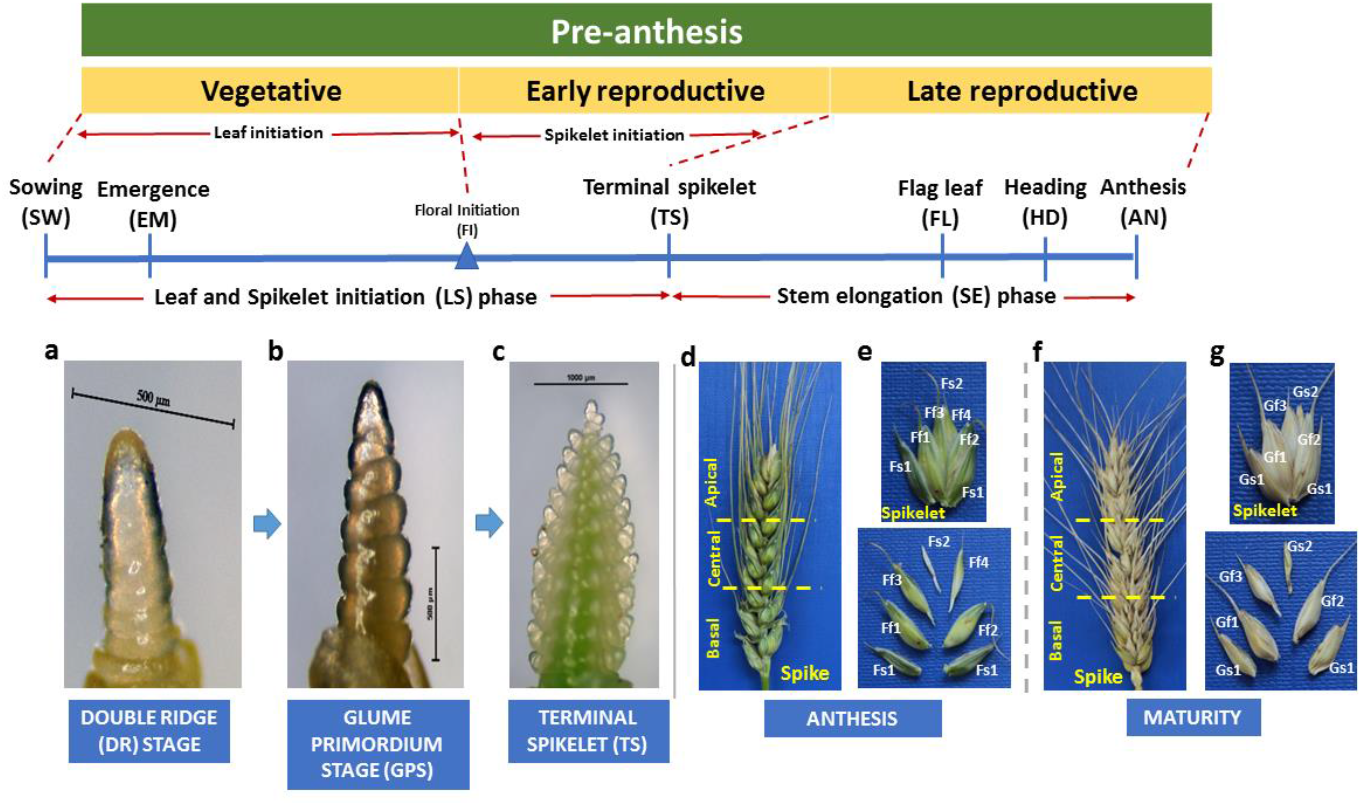
Pre-anthesis developmental phases in wheat. The vegetative phase (i.e. the leaf initiation phase) includes sowing (SW), seedling emergence (EM) and Floral initiation (FI); the early reproductive phase from FI to terminal spikelet (TS) is the phase when spikelets are initiated. In the late reproductive phase, stem elongation (SE) is subdivided into flag leaf initiation (FL), heading (HD) and anthesis (AN). The floral developmental stages during the leaf and spikelet initiation phase (LS) are represented by images of the double ridge (DR) stage (a), glume primordium stage - GPS (**b**), and terminal spikelet – TS stage (**c**). To monitor floral dynamics, spikes were dissected at both anthesis (**d**) and maturity (**f**) in three spatial levels of the spikes [basal, central (middle) and apical spikelets]. Spike dissection throughout the spikelet reveals the number of sterile (s1, s2) and fertile (f1, f2, f3, f4) florets (**e**) at anthesis or grains (**g**) at maturity.

Variation in the number of spikelets of spikes (an outcome of the spikelet initiation process taking place from floral initiation to the TS stage) may contribute to yield (Kuzay et al., 2022), in part at least due to differences in the duration of the spikelet initiation phase. Likewise, longer SE may increase fertile florets (FF) spikelet^-1^ and subsequently may support higher GY. Extended SE duration was associated with increased SDMa and consecutively increase GN and GY potential (Slafer et al., 2015). This increases the amount of accumulated resources over a particular developmental phase because the crop has more time in which it intercepts more radiation and produces higher photo-assimilates (González et al., 2011; Slafer et al., 2005). This might hint that SE duration might affect final GN via minimizing floral abortion and increasing FSR at anthesis (Borras-Gelonch et al., 2012; Miralles and Slafer, 2007). Indeed, there have been simulation studies highlighting that lengthening SE increased both grain number and yield (Hu et al., 2022). The GN spikelet^-1^ at maturity is determined by the number of FF produced during the pre-anthesis time, as well as by grain abortion that may occur soon after anthesis. Then, in post-anthesis, TKW may eventually compensate for FF aborted due to genetic and environmental factors (Ferrante et al., 2012). The reduction of floret abortion (lower number of sterile florets, SF) may be a promising breeding strategy for enhanced GN and GY potential (Sakuma et al., 2019). The association between SE duration, SDMa and GN was documented in field studies under temperate climatic regions (Borras-Gelonch et al., 2012; Guo et al., 2018), but hardly under the Mediterranean or semi-arid growing conditions of the Middle East characterized by relatively short growing season. The present study investigates whether this association is maintained for modern wheat cultivars under the short-season characteristic of the Middle East.

In the present study, a panel of nine elite Israeli bread wheat cultivars were characterized for the duration of pre-anthesis developmental phases and a number of spike fertility traits, dry matter accumulation and yield components across five Mediterranean environments in order to study: (1) the varietal differences and genotype-environment interaction (GxE) in pre-anthesis phases duration under the Mediterranean conditions, (2) whether this variability may influence biomass accumulation and partitioning and spike related traits, and (3) to what extent, pre-anthesis variability is associated with genotypic differences in yield components.

## 2. Material and methods

### 2.1. Plant materials and field experimental design

A total of nine spring bread wheat cultivars were used in the current study (from early to late phenology: ‘Yuval’, ‘Zahir’, ‘Gedera’, ‘Negev’, ‘Bar-Nir’, ‘Amit’, ‘Binyamin’, ‘Galil’ and ‘Ruta’). Field experiments were conducted under the Mediterranean environments of Central Experimental Farm (ARO, Rishon Lezion, Israel) and Gilat Research Center (ARO, Negev, Israel). The latter is more arid than the former. Three experiments were conducted in 2016-17 (season 1): early- (in early November) and late-sowing (in late December) at Gilat (Exps1 and 2, respectively), and late-sowing in late-December in Central Experimental Farm (Exp3). In each experiment, a randomized complete block design (RCBD) with four replicates was implemented. Individual plot sizes were 45, 54 and 14 m^2^ for Exp1, Exp2 and Exp3, respectively. A subset of the cultivars panel (n=6) was further evaluated in 2017-18 (season 2) at Gilat (Exp4) and Central experimental farm (Exp5), both sown at the end of November (mid-sowing) with plots (20 and 14 m^2^, respectively) again arranged in an RCBD design with four replicates.

Information for each of the 5 environments (latitude, longitude, altitude, soil types, annual rainfall and irrigation) is provided in Table 1. A sowing rate of 220 seeds m^-2^ was applied in all experiments. If required, supplementary irrigation was implemented up to anthesis to avoid the severe drought stresses during pre-anthesis development (excluding Exp5, which was exclusively rainfed during 2017-18). The experimental plots were periodically treated with fungicides and pesticides to prevent the development of any fungal pathogens or insect pests. The experimental fields in Gilat (Exp1, 2 and 4) were treated with herbicides and in the ARO central experimental farm (Exp3 and Exp5), weeds were manually removed once a month. For the complete cropping season, temperature (maximum, minimum and average), and seasonal precipitation data were collected from the online database of the Gilat-Meteorological station nearest (150 m) to each experimental field (Table S1). In the ARO central experimental farm, temperature data were recorded automatically (ten minutes intervals) by HOBO data loggers (Onset Computer Corporation, MA, USA) which were placed in the middle of the field.

**Table 1.** Details of geographical locations of studied environments and meteorological data for the season 1 (2016-17) and season 2 (2017-18)

| Season | Environment | Sites | Field name | Sowing and emergence time | Latitude (N) | Longitude (E) | Altitude (m) above sea level | Soil type | Seasonal rainfall (mm) | Supplemented irrigation (mm) | Total (mm)<sup>1</sup> |
| --- | --- | --- | --- | --- | --- | --- | --- | --- | --- | --- | --- |
| 2016-17<br>(Season 1) | Exp1 | Gilat research farm<sup>a</sup> | GS<sup>2</sup> | 09.11.2016 (Early sowing)<br>20.11.2016 (Emergence) | 31° 19' 59" N | 34° 39' 57" E | 148 m | Calcic Haploxeralf (sandy loam) | 325 | 140 | 465 |
|  | Exp2 | Gilat research farm<sup>a</sup> | GP | 22.11.2016 (Mid- sowing)<br>05.12.2016 (Emergence) | 31° 20' 33" N | 34° 40' 36" E | 161 m | Calcic Haploxeralf (sandy loam) | 292 | 108 | 400 |
|  | Exp3 | ARO central farm<sup>b</sup> | BdH | 22.12.2016 (Late sowing)<br>07.01.2017 (Emergence) | 31° 59' 38" N | 34° 49' 09" E | 40 m | Rhodoxeralf soil | 402 | 200 | 602 |
| 2017-18<br>(Season 2) | Exp4 | Gilat research farm<sup>a</sup> | GO | 16.11.2017 (Mid-sowing)<br>01.12.2017 (Emergence) | 31° 20' 2" N | 34° 40' 36" E | 153 m | Calcic Haploxeralf (sandy loam) | 215 | 80 | 295 |
|  | Exp5 | ARO central farm<sup>b</sup> | ZF | 19.11.2017 (Mid-sowing)<br>11.12.2017 (Emergence) | 32° 0' 34" N | 34° 50' 34" E | 50 m | Deep fertile vertisol | 380 | - | 380 |

<sup>1</sup> Cumulative value throughout the cropping season (sowing time-April).<sup>2</sup> GS - Gilat Sha'ar; GP - Gilat Pardes; BdH - Bet dagan Hodia; GO - Gilat Ofot; ZF - Zafariya; <sup>a</sup> Mediterranean environment which is more arid in nature <sup>b</sup> Mediterranean environment.

### 2.2. DNA extraction and genotyping with Ppd-1 and Vrn-1 genes

Genomic DNA of the studied bread wheat cultivars was extracted from fresh leaf tissue of 14-day-old seedlings following the CTAB method (Kidwell and Osborn, 1992). The germplasm panel was screened with DNA markers for *Ppd-1* (*Ppd*-A1, *Ppd*-B1, *Ppd*-D1) and *Vrn-1* (*Vrn*-A1, *Vrn*-B1, *Vrn*-D1) genes for photoperiod and vernalization sensitivity, respectively (as outlined by Curzon et al., 2019).

### 2.3. Determination of pre-anthesis phases

Pre-anthesis developmental growth phases were divided into leaf and spikelet initiation (LS) phase [includes two stages: seedling emergence (EM; from seed sowing (SW) to emergence (EM)) and terminal spikelet formation (TS; from EM to TS)] and stem elongation (SE) phase [includes three sub-phases or stages: flag leaf appearance (FL; from TS to FL), heading (HD; from FL to HD) and anthesis (AN; from HD to AN)]. All experiments were sown to wet soil or were irrigated immediately after sowing. Except for TS stage, stages were monitored in the field plots as days from the time of emergence based on 50% of experimental plot plants reaching that particular stage. In parallel, these developmental phases were recorded on a single-plant basis (three random plants collected from the middle rows of each plot) starting from the 5-leaf stage to AN. Phenological records were collected weekly (daily around the onset of TS) (Elía et al., 2016) and assigned with their respective Zadoks’s Code (ZC) (ZC10 for EM, ZC37 for FL, ZC50 for HD and ZC65 for AN) (Zadoks et al., 1974). In each plant sampled, the tillers (TL) were counted and the main stem (MS) spike was dissected and examined under a stereo microscope for the TS stage (that, depending on the GxE conditions coincided with ZCs30-33). Thermal time, in form of cumulative Growing Degree Days (GDD, °C d), was calculated for each pre-anthesis developmental phase from the seedling emergence to anthesis (SW-EM, EM-TS, TS-FL, FL-HD, HD-AN) with their respective duration in days as per the equation of Guo et al. (2018) assuming 0°C as the base temperature for wheat. Thermal time of LS, SE and total anthesis (from sowing to anthesis) time as the GDD from SW to TS (SW-TS), GDD from TS to AN (TS-SE), and GDD from SW to AN (SW-AN), respectively.

### 2.4 Plot-based phenotypic evaluation of biomass and yield components

After seedling emergence, randomly 0.5-1.0 m area from the mid plot was marked by the wooden markers which are reserved for the sampling during anthesis and maturity. At both anthesis and maturity, the sampling was done from that marked 0.5 m (for Exp1, Exp2 and Exp3) or 1 m (for Exp4 and Exp5) in the middle row of each plot (different sample size length depending on total plot size at each experimental site) followed by the procedure of Elía et al. (2016) and Terrile et al. (2017). This is to analyse the major anthesis and maturity data based on the plants in equal size of the marked area having correct density and emergence time across all the experimental plots. At anthesis, plants were separated into MS and TL to count their number as well as the number of spikes; and then to determine AGDM and SDMa following the protocol used by Elía et al. (2016) and Terrile et al. (2017). The same process was repeated at maturity for the above-mentioned traits. Fruiting efficiency (FE) was estimated as a ratio between grain number at maturity and spike dry matter (SDMa) as per Elía et al. (2016). Spikes of MS and TL were dried, weighed and then threshed. The yield components were measured by counting grains in a machine (Model S-25; Data technologies Ltd., Jerusalem, Israel) and weighing them. PH was measured from the ground to the tip of MS spike (excluding awns).

### 2.5. Spike trait characterization and phenotypic evaluation

Three random plants were sampled at both anthesis and maturity in each plot. Each plant was further separated into MS and TL and each spike was measured for DM partitioning and floret conditions. The number of fertile florets (FF, florets with all the structural and reproductive organs are well developed) and sterile florets (SF, floret positions in that the structural organs, rachilla and glumelles, are well developed but the reproductive organs of the floret primordium, androecium and gynoecium, were died) were counted in each of the spikelet positions considered [basal (3^rd^-4^th^ spikelets most proximal to the peduncle), central (spikelet located in the middle of the spike) and apical (3^rd^-4^th^ spikelet from the distal spike part)], followed by oven drying for DM determination of all spike components as described in Elía et al. (2016). Besides, the number of FF and SF spikelet^-1^ (indexed by its position along with the rachis node), the total number of spikelets spike^-1^, the total number of florets or TF (FF+SF) spike^-1^ and floret survival rate (FSR%, i.e. the proportion of FF respect to TF) of the spike were determined in three different spike levels separately as mentioned above. Individual spike dry matter (SDM) was measured at both anthesis (SDMa) and maturity (SDMm). SDMa includes the biomass of rachis (R), FF and SF; whereas SDMm includes the biomass of chaff (rachis, glumes, lemma and palea) and grains (i.e. the grain weight, GW). Also, at both anthesis and maturity, total above-ground dry matter (AGDM) was measured in this subsample as done for the whole sample, which includes the aboveground vegetative (stem and leaf) and reproductive (spike) biomass. The data taken based on the single plant basis were upscaled to the whole sample considering the ratio of biomass in these sub-sampled plants and that of the whole sample.

### 2.6. Statistical analysis

The JMP^®^ Pro version 14.0 statistical package (SAS Institute Inc., North Carolina, USA) was used for all statistical analyses. Descriptive statistics were performed on the full dataset to illustrate variable distribution. ANOVA was carried out considering the genotype and environment as a fixed effect and the block as a random effect. Pearson correlation analyses and linear regression model were applied to visualize the association relationship of pre-anthesis developmental phases with biomass accumulation, floret dynamics and yield components. Principal Component Analysis (PCA) was performed for each season (season 1 and season 2) based on the mean values for each variety in a specific environment. Highly auto-correlated variables and values with low loading scores were removed from the analysis. Contrast analysis has been applied to statistically compare the means of various traits at both anthesis and maturity between the selected cultivar pairs with significant differences in TS and SE phase with no significant differences in total time to anthesis at a specific environment.

## 3. Results

### 3.1. Genotypic characterization for PPD-1 and VRN-1 genes

In general, very low polymorphism was found in the allelic profile of the wheat cultivars (Table 2). All nine cultivars in the panel were found to carry the photoperiod sensitive (SEN) alleles in the A and B genomes, *Ppd-A1b* and *Ppd-B1b*, respectively. However, they were largely insensitive to photoperiod due to the presence of the dominant photoperiod insensitive (INS) allele (*Ppd-D1a*) in the D1 genome. On the other hand, higher allelic variability was detected in the *Vrn* loci of the three genomes, but in all cultivars, there were more insensitive (*Vrn1*, conferring spring habit) than sensitive (*vrn1*, conferring winter habit) alleles.

**Table 2.** Genotypic characterization of allelic profiling of *PPD1* (photoperiod) and *VRN* (vernalization) genes in the tested panel of modern Israeli bread wheat cultivars

| Cultivars | Photoperiod |  |  | Vernalization |  |  |
| --- | --- | --- | --- | --- | --- | --- |
|  | *PPD-A1* | *PPD-B1* | *PPD-D1* | *VRN-A1* | *VRN-B1* | *VRN-D1* |
| Yuval | *Ppd-A1b*<br>(SEN<sup>1</sup>) | *Ppd-B1b*<br>(SEN) | *Ppd-D1a*<br>(INS<sup>1</sup>) | *Vrn-A1*<br>(INS) | *Vrn-B1*<br>(INS) | *Vrn-D1*<br>(INS) |
| Zahir | *Ppd-A1b*<br>(SEN) | *Ppd-B1b*<br>(SEN) | *Ppd-D1a*<br>(INS) | *vrn-A1*<br>(SEN) | *Vrn-B1*<br>(INS) | *Vrn-D1*<br>(INS) |
| Gedera | *Ppd-A1b*<br>(SEN) | *Ppd-B1b*<br>(SEN) | *Ppd-D1a*<br>(INS) | *vrn-A1*<br>(SEN) | *Vrn-B1*<br>(INS) | *Vrn-D1*<br>(INS) |
| Negev | *Ppd-A1b*<br>(SEN) | *Ppd-B1b*<br>(SEN) | *Ppd-D1a*<br>(INS) | *vrn-A1*<br>(SEN) | *Vrn-B1*<br>(INS) | *Vrn-D1*<br>(INS) |
| Bar-Nir | *Ppd-A1b*<br>(SEN) | *Ppd-B1b*<br>(SEN) | *Ppd-D1a*<br>(INS) | *Vrn-A1*<br>(INS) | *Vrn-B1*<br>(INS) | *vrn-D1*<br>(SEN) |
| Amit | *Ppd-A1b*<br>(SEN) | *Ppd-B1b*<br>(SEN) | *Ppd-D1a*<br>(INS) | *vrn-A1*<br>(SEN) | *Vrn-B1*<br>(INS) | *Vrn-D1*<br>(INS) |
| Binyamin | *Ppd-A1b*<br>(SEN) | *Ppd-B1b*<br>(SEN) | *Ppd-D1a*<br>(INS) | *Vrn-A1*<br>(INS) | *Vrn-B1*<br>(INS) | *vrn-D1*<br>(SEN) |
| Galil | *Ppd-A1b*<br>(SEN) | *Ppd-B1b*<br>(SEN) | *Ppd-D1a*<br>(INS) | *vrn-A1*<br>(SEN) | *Vrn-B1*<br>(INS) | *Vrn-D1*<br>(INS) |
| Ruta | *Ppd-A1b*<br>(SEN) | *Ppd-B1b*<br>(SEN) | *Ppd-D1a*<br>(INS) | *vrn-A1*<br>(SEN) | *Vrn-B1*<br>(INS) | *Vrn-D1*<br>(INS) |

<sup>1</sup>SEN - Sensitive; INS – Insensitive

### 3.2. Pre-anthesis developmental phases

There was variability in time to anthesis (Figure 2). Yuval was the earliest flowering cultivar across experiments, only in Exp1 there was a shorter cultivar (Zahir), although this difference in earliness was not statistically significant (Figure 2a), and Zahir was consistently the second earliest cultivar across the other four environments (Figure 2). In season 1, Ruta (only grown in the first season) was the latest flowering cultivar, followed by Galil and Negev (with confirmed lateness in the second growing season; Figure 2). There was also variability in the partitioning of the time to anthesis into its two major sub-phases. We identified all possible combinations of cultivars with a similar time to anthesis, but significantly different TS and/or SE durations (Figure 2).

**Figure 2.**
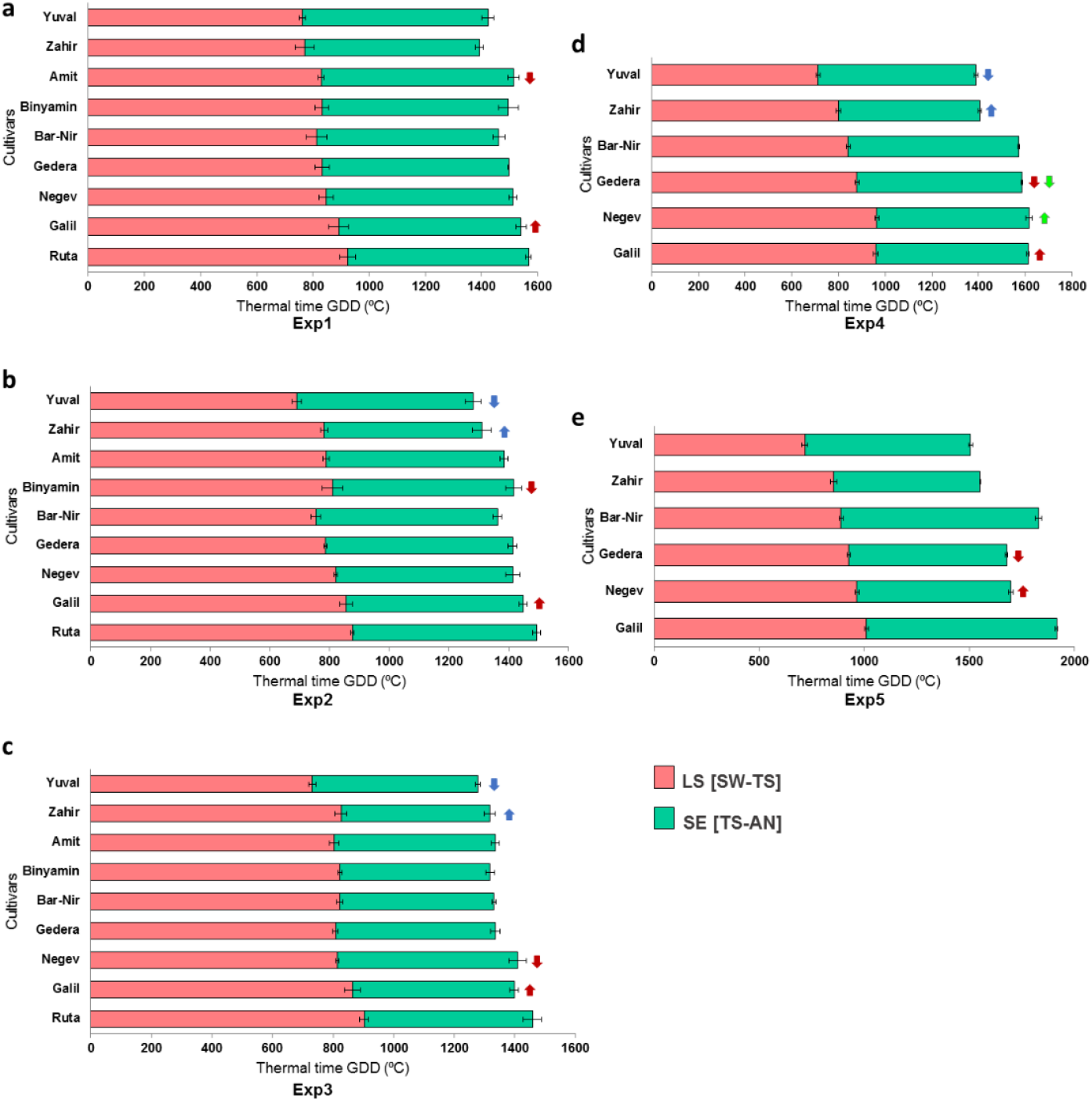
Pre-anthesis developmental phases of a panel of modern bread wheat cultivars. Phenology of pre-anthesis developmental phases in two seasons (2016-17 and 2017-18) across five environments: Exp1 (**a**), Exp2 (**b**), Exp3 (**c**), Exp4 (**d**), and Exp5 (**e**). Developmental phases were considered and delimited by the following stages: sowing (SW), seedling emergence (EM, ZC10), terminal spikelet formation (TS, ZC31), flag leaf stage (FL, ZC37), heading (HD, ZC50) and anthesis (AN, ZC65). Here we represent the leaf and spikelet (LS) initiation phase from SW to TS, and stem elongation (SE) phase from TS to AN. Colour arrows within environments represent pairs of cultivars whose thermal time from SW to AN was similar (non-significant difference), but significantly differed in duration for SW-TS and TS-AN in opposite directions [longer SW-TS and shorter TS-AN marked with an upward arrow (↑), and shorter SW-TS with longer TS-AN with a downward arrow (↓); the colour of the arrows identify the particular pairs].

Generally, in all cultivars across environments, LS phase - the duration of SW-TS, was longer than SE i.e., TS-AN. When focusing on the genotypic variation, differences in time to anthesis within each environment were better related to those in SW-TS (Figure 3a) than to those in the SE phase (Figure 3b).

**Figure 3.**
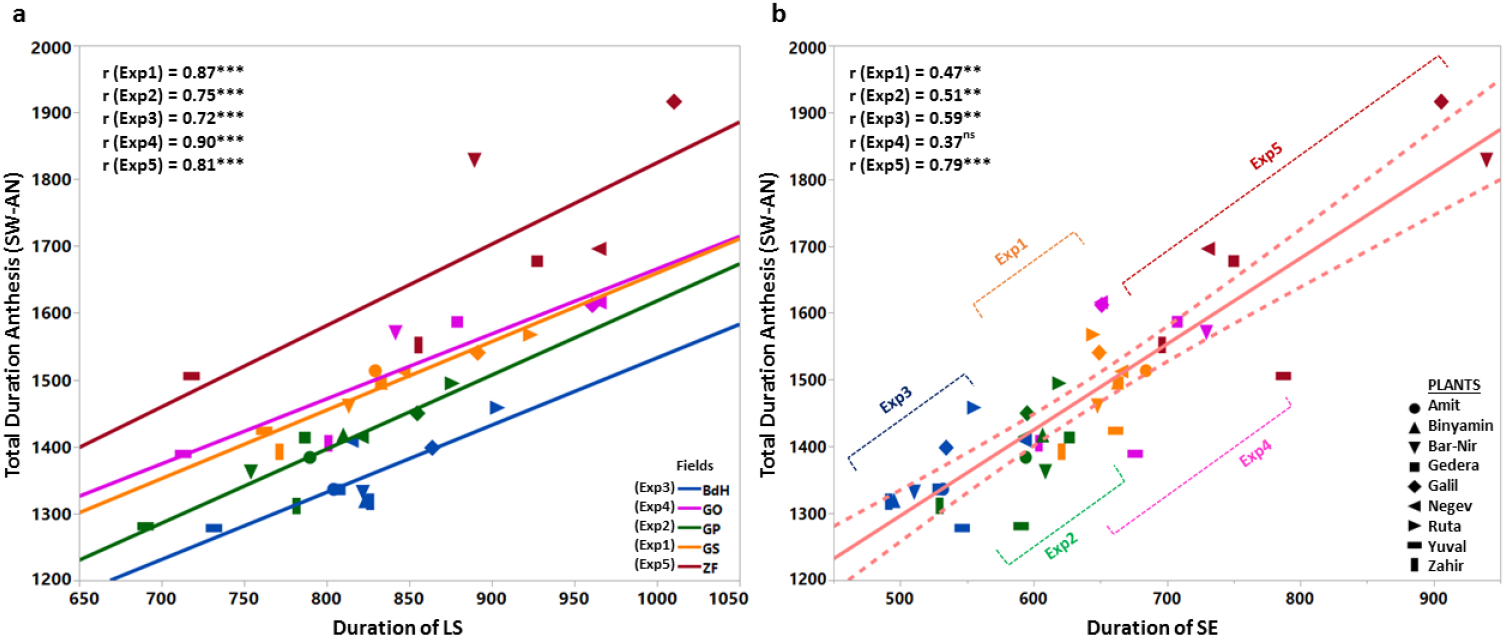
**Linear regression of total anthesis time (SW-AN) to LS (a) and SE (b)** durations for modern bread wheat cultivars across five environments (Exp1-5). The mean values of each cultivar for LS, SE and total anthesis duration were plotted for each environment. The correlation coefficient (r) and significance level (p≤0.001) for cultivars under each environment are presented in the graph inset. Different environments are marked in colours and different marker symbols represent different cultivars (see the legend).

### 3.3. Association of LS and SE with spike traits, dry matter and yield-related traits

Both the LS and SE duration were significantly associated with several spike traits, dry matter and yield components. However, no significant association was observed between SE and other traits in Exp1 and Exp2, which highlights the LS associations as a bit more prominent. For example, LS duration was positively correlated with anthesis traits like spikelets spike^-1^, SF spike^-1^ in Exp1; FF spike^-1^ and spikelet^-1^, FSR in Exp2; FFDM, SDMa spike^-1^, FSR spike^-1^ in Exp3; and the maturity traits like GN spike^-1^, GN m^-2^, GW spike^-1^, GW m^-2^, CW spike^-1^ and AGDM in Exp4 (Table 3; S2). The duration of major phases of pre-anthesis (EM-TS, TS-AN, EM-AN) are somewhat associated with FF number, biomass partitioning (AGDM and SDMa) and FSR at anthesis and FF number, grain yield (GN) and FE across the five environments, but the results were not static in every case. No significant associations have been found in TS-AN phase and also for Env3 (Table 3). The LS duration also negatively correlated with TKW in both Exp1 and Exp2. Similarly, SE duration was positively correlated only with the majority of anthesis parameters in Exp3 like FF spike^-1^, SF spike^-1^ and TF spike^-1^ and two of the maturity traits: GN spike^-1^ and PH in Exp5. For the major anthesis traits and yield components, SE was negatively associated with FFDM, SDMa spike^-1^, SF spike^-1^, GW spike^-1^, GW m^-2^ and TKW in Exp4 (Table S2). However, for all experiments with moderate or no water stress (all those receiving supplementary irrigation), any effects of EM-TS on yield components produced compensations so that neither the duration of EM-TS or TS-AN affected yield significantly nor grains per m^2^. The exception was Exp4, the one with more severe water stress, where the duration of SW-TS, responsible for the total time to AN, was positively related to yield and grains per m^2^ (Table S2).

**Table 3.** Correlation co-efficient (r) for pre-anthesis phases with major anthesis and maturity traits across five environments.

| Traits | Pre-anthesis phases | Environments |  |  |  |  |
| --- | --- | --- | --- | --- | --- | --- |
|  |  | Exp1 | Exp2 | Exp3 | Exp4 | Exp5 |
| FFa spike<sup>-1</sup> | EM-TS | 0.36* | 0.42* | 0.54*** | -0.04 | 0.67** |
|  | TS-AN | -0.01 | -0.02 | 0.40* | -0.29 | 0.10 |
|  | EM-AN | 0.31 | 0.36* | 0.72*** | -0.17 | 0.49* |
| AGDM | EM-TS<sup>1</sup> | 0.51 | 0.23 | 0.15 | 0.83* | 0.88* |
|  | TS-AN | -0.19 | -0.11 | 0.37 | 0.18 | 0.34 |
|  | EM-AN | 0.41 | 0.14 | 0.33 | 0.85* | 0.78 |
| SDMa spike<sup>-1</sup> | EM-TS | 0.19 | -0.10 | 0.33* | 0.57** | 0.80*** |
|  | TS-AN | 0.07 | -0.24 | 0.11 | -0.09 | 0.46* |
|  | EM-AN | 0.20 | -0.25 | 0.35* | 0.48* | 0.79*** |
| FSR | EM-TS | 0.32 | 0.42* | 0.37* | 0.02 | 0.75*** |
|  | TS-AN | -0.02 | -0.08 | -0.07 | -0.35 | 0.29 |
|  | EM-AN | 0.27 | 0.32 | 0.25 | -0.15 | 0.66** |
| FFm spike<sup>-1</sup> | EM-TS | 0.30 | 0.23 | 0.14 | 0.29 | 0.26 |
|  | TS-AN | 0.02 | 0.36* | 0.37* | -0.18 | 0.38 |
|  | EM-AN | 0.28 | 0.45** | 0.38* | 0.18 | 0.40 |
| GN m<sup>-2</sup> | EM-TS | 0.04 | 0.30 | -0.11 | 0.76*** | -0.01 |
|  | TS-AN | 0.21 | 0.29 | -0.18 | -0.22 | -0.26 |
|  | EM-AN | 0.14 | 0.45** | -0.22 | 0.58** | -0.16 |
| GN spike<sup>-1</sup> | EM-TS | 0.28 | 0.51** | 0.12 | 0.73*** | 0.45 |
|  | TS-AN | 0.12 | 0.05 | -0.12 | -0.37 | 0.46* |
|  | EM-AN | 0.31 | 0.48** | 0.01 | 0.49* | 0.57* |
| FE | EM-TS | 0.81** | 0.73* | -0.08 | 0.53 | -0.94** |
|  | TS-AN | -0.21 | 0.52 | -0.14 | 0.52 | -0.30 |
|  | EM-AN | 0.68* | 0.82** | -0.14 | 0.73 | -0.80 |

<sup>1</sup>EM-TS = from Emergence to TS; TS-AN = from TS to Anthesis; EM-AN = from Emergence to Anthesis; FFa spike<sup>-1</sup> = Number of fertile floret florets per spike at anthesis; AGDM = above ground dry matter m<sup>-2</sup> at anthesis; SDMa spike<sup>-1</sup> = Spike dry matter per spike at anthesis; FSR = floret survival rate (%) at anthesis; FFm spike<sup>-1</sup> = Number of fertile floret florets per spike at maturity; GN m<sup>-2</sup> = grain number per 1m<sup>2</sup> area at maturity; GN spike<sup>-1</sup> = grain number per spike at maturity; FE = fruiting efficiency in 1m during maturity. '\*', '\*\*' and '\*\*\*' denote the significance level at p≤0.05, p≤0.01 and p≤0.001, respectively.

### 3.4. Phenotypic multivariate analyses across environments and genotypes

ANOVA of the duration of pre-anthesis phases, spike traits at anthesis and maturity and yield components were implemented only for the six cultivars which were grown in all five environments and showed significant genotype-environment interaction (GxE) for the studied traits (Table S3). PCA was performed combinedly with six cultivars in five environments. Similarly, to the ANOVA (Table S3), PCA charts exhibit a clear separation of environments and to a lesser extent cultivars within each environment. In Figure 4, the variety panel is plotted with PC1 explaining 42.8% of the phenotypic variation and was positively loaded with three spike traits at anthesis (spikelet spike^-1^, FF spike^-1^ and TF spike^-1^), LS duration and three yield-related traits (TF spike^-1^, GN spike^-1^ and GW spike^-1^) measured at maturity. PC1 was also negatively loaded with the total number of tillers m^-2^ at maturity. PC2 explained 26.2% of the variation in the dataset and was positively loaded with three DM-associated traits measured at anthesis (total plants m^-2^, SDMa m^-2^ and AGDM m^-2^), SE duration and one yield-associated trait (HI) at maturity. All the major traits at both anthesis and maturity were loaded positively, except the total number of mature TL. In the PCA plot, the five environments (Exp1, Exp2, Exp3, Exp4 and Exp5) form distinctive clusters across the phenotypic space (each encircled by dotted lines in different colours). Thus, in the variation of GxE, the effect of environment was prevalent that of the genotypes (Fig. 4a; Table S3). The late-sowing environment (Exp3) is widely distributed in both PC axes located mainly in the negative quarter of the PCA chart which is characterized by low productivity of all the traits. Early-sowing (Exp1) environment is characterized by the high productivity in spike traits at both anthesis and maturity, and it is distributed positively on PC1 (Figure 4a). In each environment, crop phenology was the dominant discriminator between the cultivars, which were all framed between the boundaries of Yuval (early) and Galil (late). This is also supported by the single Pearson-correlation analysis. This is evident from Figure 4b, LS and SE duration was individually significantly associated with major spike traits (spikelet number spike^-1^, FF and TF spike^-1^), DM (SDM, AGDM m^-2^) and yield components (GN spike^-1^, GW spike^-1^, GN m^-2^, GW m^-2^, TKW) (Table S2). Although LS and SE duration are positively related (Figure 4b), it mostly reflects the environmental effects as late sowing (Exp3) would have reduced the length of both pre-anthesis phases (Figure 4a). Exp4 was characterized by lower productivity in spike and yield-related traits at both anthesis and maturity compared to Exp5. Under all environmental conditions, the cultivars with long LS phase (Zahir, Negev and Galil) were positively loaded relative to their short LS cultivar counterpart (Yuval, Gedera and Bar-Nir).

**Figure 4.**
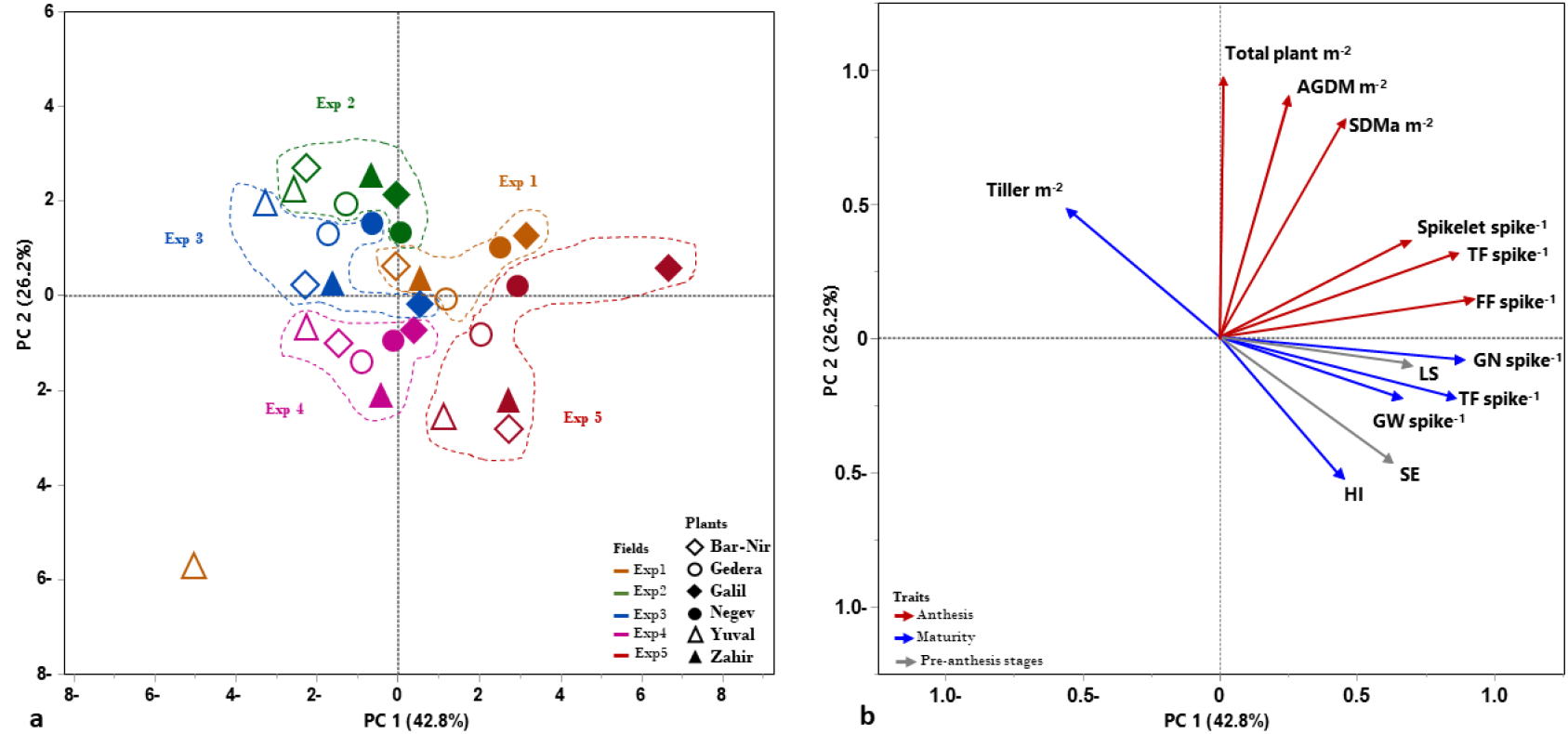
**Principal component analysis (PCA) of spike traits at anthesis and maturity and yield components of** six wheat cultivars in five environments (Exp1, Exp2, Exp3, Exp4 and Exp5) across both Season 1 (2016-17) and Season 2 (2017-18) (**a**) for the studied traits (**b**). The timing of measurement of traits at anthesis (RED) or maturity (BLUE) is marked. PC factor loadings are expressed as vectors (arrows) using the coordinates for PC1 and PC2. Individual cultivar’s PC scores are marked by a dotted coloured line to define their common environment. Cultivars are Yuval, Zahir, Gedera, Negev, Galil and Bar-Nir are marked with different symbols (see legend). The traits are Total plant m^-2^ (total number of plants in 1m^2^ area), Tiller m^-2^ (total number of tillers in 1m^2^ area), SDMa m^-2^ (total spike dry matter at anthesis in 1m^2^ area), AGDM m^-2^ (above ground dry matter in 1m^2^ area), FF spike^-1^ (number of fertile florets per spike), TF spike^-1^ (total number of florets per spike), spikelets spike^-1^ (total number of spikelets per spike), LS duration, SE duration, GN spike^-1^ (grain number per spike), GW spike^-1^ (grain weight per spike) and HI (harvest index).

### 3.5. Association of pre-anthesis phases with floral conditions and spike fertility

Comparative analyses of spike fertility, considering the number of fertile and sterile florets in each of the spikelets, were performed across all environments for early and intermediate pairs of cultivars ‘Zahir-Yuval’ (Figure 5) and ‘Negev-Gedera’ (Figure S1). Within each pair, EM-TS and TS-AN duration was significantly different, although anthesis duration (EM-AN) was similar. The mean comparison of FSR of the central spikelets has clearly illustrated that those cultivars with longer SE are significantly superior (except ‘Negev-Gedera’ at Exp2, Figure S1b3). Compared to shorter LS cultivars (Yuval and Gedera), longer LS cultivars (Zahir and Negev) have higher mean FSR (%) and grain set % at both anthesis and maturity, respectively. In general, the total number of spikelets spike^-1^ was significantly higher in the cultivars with longer LS (Zahir and Negev) when compared to their respective shorter SE pairs (Yuval and Gedera) at both anthesis and maturity, in most of the environments (Figure 5 a4, b4, d4, e4; Figure S1 a4, e4).

**Figure 5.**
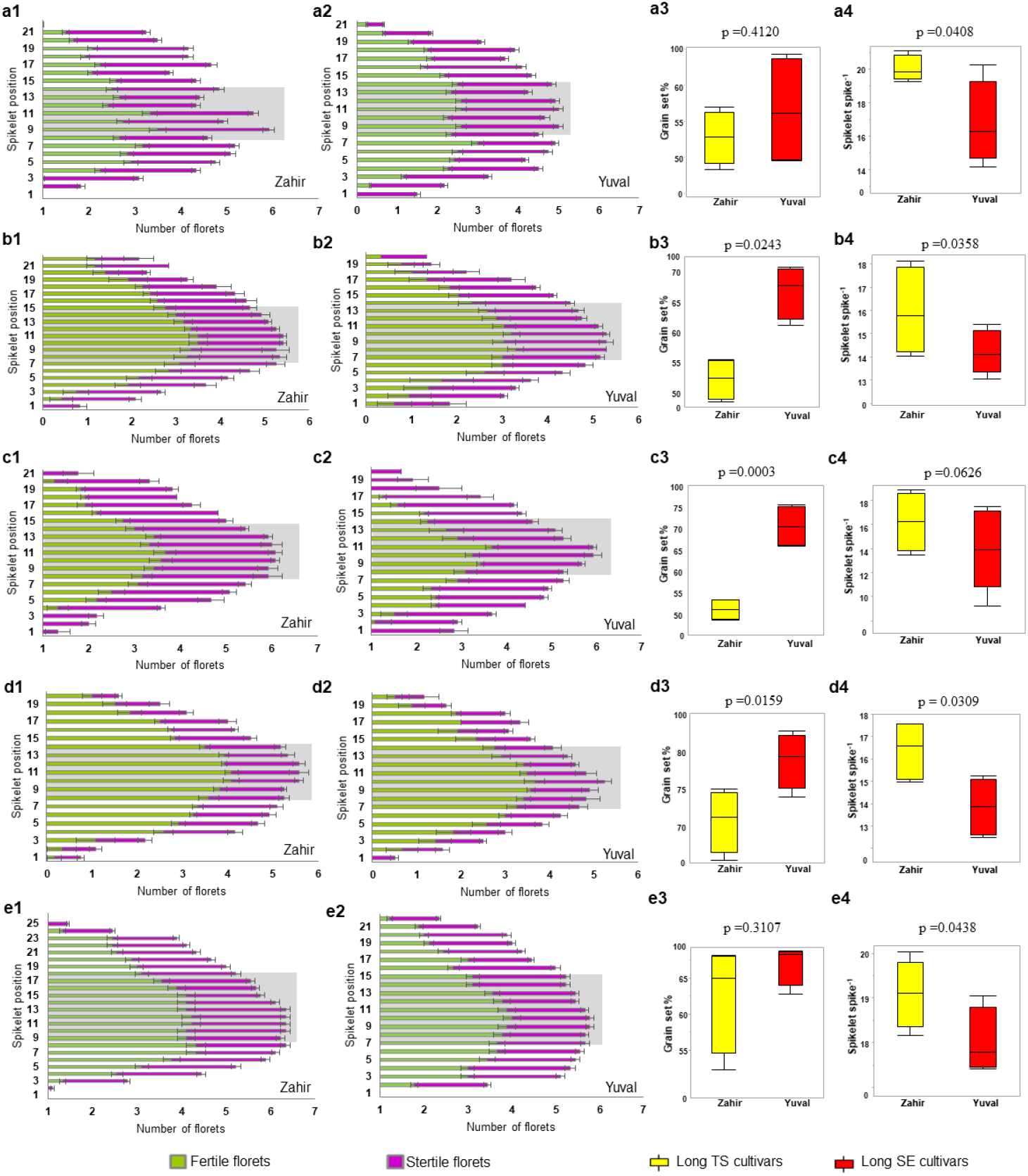
Floret conditions in the spike between the cultivar pair - Zahir and Yuval. The number of grains (results of fertile florets at anthesis) and sterile florets of the cultivar pairs are representing as the bar graph according to each spikelet position from the basal to apical part of the rachis node of each main-shoot spike at maturity. **a** represents Exp1, **b** represents Exp2, **c** represents Exp3, **d** represents Exp4, and **e** represents Exp5. The Grey shaded part represents the position of central spikelets. The boxplots are representing the range of grain set % of the central spikelets at maturity (a3, b3, c3, d3, e3) and the mean spikelet number spike^-1^ including both main-shoot and tillers at maturity (a4, b4, c4, d4, e4) of Zahir-Yuval pair in all five environments. The ‘*t*’ values for the significance probability between the pair are given in each plot.

### 3.6. Association of pre-anthesis phases with spike traits at anthesis and yield components at maturity

Contrast analyses detected significant differences within different cultivar pairs for nine spike traits and dry matter at anthesis *viz*. reproductive organ DM (g) spike^-1^, FFDM (g) spike^-1^, spikelet numbers spike^-1^, FF spike^-1^, SF spike^-1^, TF spike^-1^, SDMa spike^-1^, SDMa m^-2^ and AGDM (g) m^-2^. (Figure 6 a-c; Figure S2). Field experiment Exp2 was an exception with no significant differences in spike traits within the cultivar pairs. Longer LS duration was associated with the increased value of the spike traits and dry matter at anthesis, except for SF spike^-1^ which was negatively associated with LS. At maturity, contrast analyses between the cultivars composing the pairs show that except for the terminal drought-prone environment (Exp5), Exp1-Exp4 environments expressed significantly higher values in the cultivar with longer LS in six major yield components at maturity *viz*. GN spike^-1^, GN m^-2^, GW spike^-1^, GW m^-2^, TKW and FSR%, within the cultivar pairs (Zahir-Yuval, Negev-Gedera, Galil-Binyamin, Galil-Amit and Galil-Negev) (Figure 6 d-f; Figure S3).

**Figure 6.**
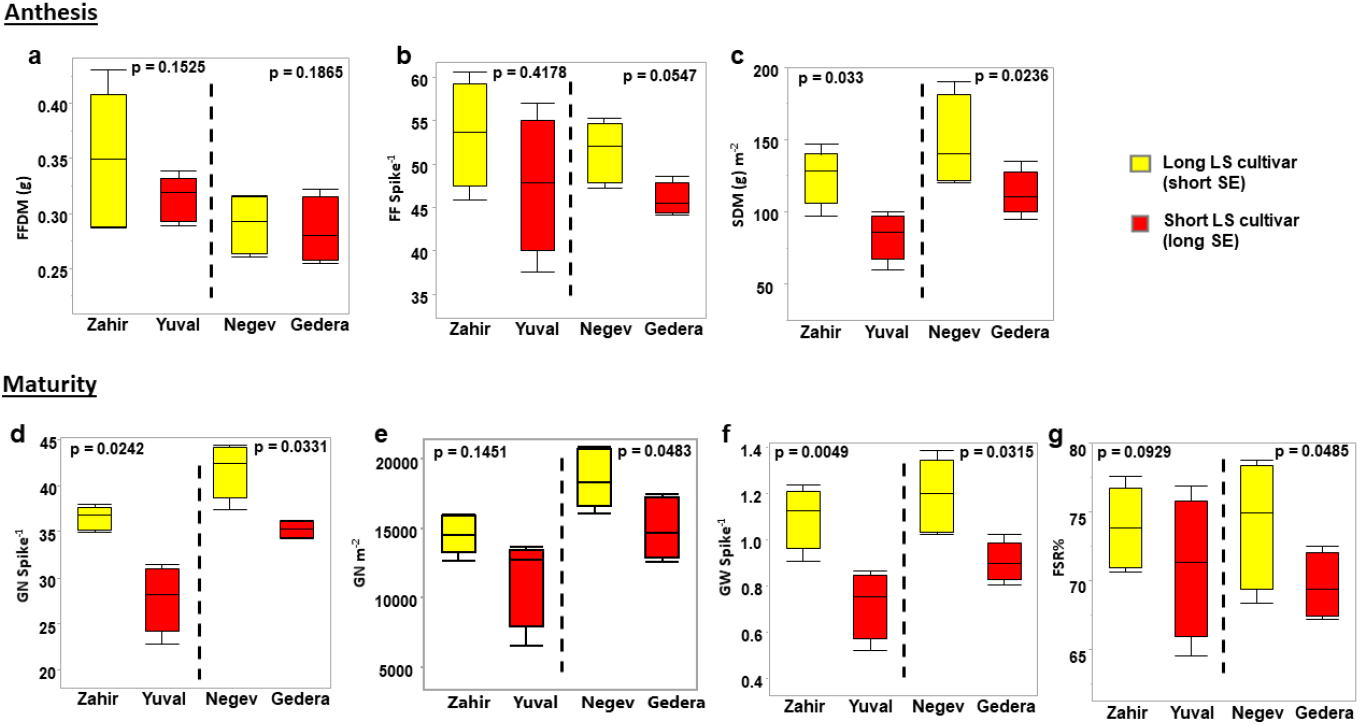
**Contrast analysis of spike traits at anthesis (a-c) and major yield components at maturity (d-g)** between the cultivar pairs (‘Zahir-Yuval’ and ‘Negev-Gedera’) in mid-sowing Exp4 (Gilat Ofot) environment, which showed significant differences in LS with non-significant anthesis time. Box plot representing (**a**) fertile floret dry matter spike^-1^, (**b**) fertile florets spike^-1^, (**c**) spike dry matter m^-2^, (**d**) grain number spike^-1^, (**e**) grain number m^-2^, (**f**) grain weight spike^-1^, and (**g**) floret survival rate (FSR%). The ‘t’ values for the significance probability between the pair are given in each plot if its value is less than or equal to 0.05, and it is non-significant if the value is beyond 0.05.

## 4. Discussions

The wheat pre-anthesis phase of SE, during which floret development takes place, was shown to be associated with DM allocation to the juvenile developing spike with a possible promoting effect on spike fertility (Elía et al., 2016; González et al., 2005, 2011; Slafer et al., 1990; Terrile et al., 2017). In the present study, we aimed to test whether this association could be extended to wheat grown under Mediterranean conditions dictating a rather short growing season. We attempted to assess the relative contribution of the duration of the LS and SE phases to spike development, DM accumulation and yield components.

The phenological amplitude (the difference between the earliest and late variety in a certain environment) of the highly adapted elite modern wheat panel was rather narrow [i.e. 175 °Cd and 158 °Cd for early (Exp1) and late (Exp3) sowing environments, respectively (Figure 2)] and well associated with the similar genetic profile of both *PPD-1* and *VRN-1* genes for photoperiod and vernalization response, respectively. All nine cultivars in the panel were found to carry photoperiod insensitive allele *Ppd-D1a* on chromosome 2D. As *PPD-D1* is most frequently the strongest photoperiod sensitivity gene (e.g. Ochagavia et al., 2017), the presence of its dominant allele (insensitive) largely masked the photoperiod sensitivity that would have been expected from the recessive alleles on the 2A (*Ppd-A1b*) or 2B (*Ppd-B1b*) chromosomes (resulting in all cultivars expressing photoperiod-insensitive). On the other hand, higher allelic variability was detected in the three *VRN* loci (*VRN-A1, VRN-B1* and *VRN-D1*). However, polymorphism of dominant (*Vrn*) and recessive (*vrn*) alleles among the panel was not phenotypically expressed, likely because no genotype had all three recessive alleles and therefore the requirements of vernalization had been relatively small and satisfied by the mild winter in the studied field locations. Chances are that most differences in phenology in this panel may be due to differences in earliness *per se* (*Eps*) genes, not genotyped here, as was already documented (e.g. Ochagavia et al., 2018).

Of course, the environment is a major factor dictating wheat crop phenology and performances as shown for the studied traits by the ANOVA results in Table S3 and by the strong environmental discrimination at single (Figure 3a and 3b) and multivariate analyses (Figure 4). As it had been already shown by Shiff et al. (2021), delay in seed sowing leads to reduced SE (i.e. Exp3 compared to Exp1-2 Figure 3b), which in turn decreases productivity, as biomass accumulation is highly dependent on the thermal timespan (plotted on the negative quarter of the PCA chart Figure 4a-b). In Mediterranean environments of the Middle East (as well as those in North Africa), the wheat growing cycle is short (i.e. less than 12 weeks from emergence to anthesis). Time to anthesis was more strongly to the duration of LS than to SE across the five environments for the studied panel, indicating that the initiation of stem elongation (phenotypically observed by the 1 detectable node above the soil surface) is a useful phenological marker for earliness in these conditions (Figure 3a).

Potential wheat GN may be dependent on the LS and SE phases. The former is responsible for the formation of both the canopy structure (responsible in turn for capturing resources) and spikelet spike^-1^ (in terms of the sink side) while the SE contributes to the formation of floral meristems in each spikelet which is later manifested as florets spikelet^-1^ (see Figure 1; Li et al., 2019). Overall, in this study, a more prominent effect was encountered for LS, rather than SE when testing the majority of anthesis traits and associated yield attributes at maturity. Altogether GN and GW, either per m^2^ or per spike, were significantly positively correlated with LS duration in three more arid fields at Gilat (Exp1, Exp2 and Exp4). However, the well-documented tradeoffs of TKW in Exp1 and Exp2 (Acreche and Slafer, 2006) prevented its expression of the LS duration in the final GY (Table S2). Spike dissection traits at anthesis showed, in general, significant positive associations between them and the FSR (Table S2). This might indicate that FF number and their DM accumulation directly improve the FSR. The strong association (r=0.78 in Exp2) between spikelets spike^-1^ and FF showed that the spikelet number promotes the number of FF and jointly supports the TF spike^-1^, at least in conditions with a limited duration of spikelet initiation.

Langer and Hanif (1973), and Sibony and Pinthus (1988) found that bread wheat floret developmental rate varies along the rachis (when comparing basal, central and apical spikelets of the spike). Central spikelets are more fertile than the other two parts bearing more florets which resulted in higher GNs at maturity in the present study (Figure 5 a-e1, a-e2; Figure S1). It was evident that longer SE cultivars had greater grain set % in central spikelets (Figure 5 a3, b3, c3, d3, e3; Figure S1 a3, b3, c3, d3, e3) and were consistent in most of the mid-sowing environments (Exp2, Exp4, Exp5). A similar trend has been reported for the longer SE associated with increased FSR at anthesis (Borras-Gelonch et al., 2012; Miralles and Slafer, 2007). However, contrary to this trend, the number of FF spike^-1^ (also FSR of the whole spike) and the spikelets spike^-1^ were consistently higher in cultivars with longer LS (Figure 5 a4, b4, c4, d4 and e4; Figure S4 a4, b4, c4, d4 and e4). This apparent contradiction might be explained by the negative association (tradeoffs) of central FSR to peripheral spike organs in both basal and apical. It is therefore important to highlight that under the short wheat-growing season, LS duration seems a key driver of SDMa, GN spike^-1^ and GW spike^-1^. TS is the key transition from LS to SE (Figure 1), which also terminate the spikelet primordia formation. Therefore, assuming no delay in heading time, LS extension or delay allows additional spikelet infrastructure buildup to enhance GN potential. Maintaining a high FF ratio at anthesis either per spikelet or per whole spike can drive higher GN at maturity, which is in the agreement with Terrile et al. (2017). Miralles and Slafer (1995) and Miralles et al. (1998) found a strong quantitative relationship of GN spike^-1^ with GN spikelet^-1^, rather than spikelet numbers spike^-1^, because of the association between GN spikelet^-1^ with FSR or the higher number of FF at anthesis.

Many reports highlighted the SE phase as the main potential contributor to GY through increasing spike DM at anthesis (Elía et al., 2016; Ferrante et al., 2012; González et al., 2011; Slafer et al., 2014; Slafer et al., 2015). Alternatively, under the studied conditions, the results highlight the potential of the LS duration in promoting biomass allocation to the spike, aboveground dry matter accumulation and increased yield components in bread wheat (Figures 6, S1 and S2). Although the last part of the SE phase (from flag leaf initiation to heading) is most coincidental with maximal spike and peduncle growth, the floret fertility (FSR) and subsequently the GN at maturity were largely pre-determined in the floral primordia stage during the LS phase.. The hypothesis of the advantage of a longer SE (Slafer et al., 2021) is mostly valid when there are no penalties in resource capture at the onset of SE due to either a sufficiently long LS or the reduced LS was compensated by increased plant density. Otherwise reducing LS would bring about the penalties in source strength (Hu et al. 2022) during the SE phase and that lack of source strength in this critical period would bring about a limited allocation of resources to the juvenile spikes reducing consequently SDMa and spike fertility (Reynolds et al., 2022). Extended SE showed a positive effect on FSR% in the central spikelets of the spike at maturity but did not improve overall spike fertility and yield components. As suggested by Sakuma et al. (2019), the reduction of floret abortion in form of the number of SF would be a promising breeding strategy for enhanced GN and yield potential in parallel to improve SDM accumulation, and lengthening the SE phase might be a tool to achieve this provided there are no penalties in reducing the length of the LS phase (as it seems to be the case in most wheat growing regions in which the crop cycle sufficiently long).

## 5. Conclusion

Under the Mediterranean and semi-arid climate, and likely in any environment characterised by having a short growing season, the LS phase duration had a stronger effect than SE, on DM accumulation in the spike at anthesis and spike GN and GW at maturity. In such a short growing cycle, a short LS phase would likely penalize the formation of source structures, which are essential for sink determination. Therefore, the hypothesis of lengthening SE as a way to improve GN and yield seems not to apply to short growing seasons (Hu et al., 2022). In this condition, the advantage of a longer SE phase at the expense of a shorter LS would only apply if the penalties could be avoided through changes in agronomic field management (e.g., increasing the sowing rate); otherwise, lengthening the SE phase would not be a valid strategy to increasing GN and wheat yield. So, the importance of the LS phase is derived from the development of an appropriate number of floret primordia and it would be very helpful to improve the final GY at maturity (Sakuma et al., 2019). The varietal combination with variable LS and SE duration could be implemented in the breeding pipeline and used as pre-breeding materials for GN improvement. Furthermore, the findings will improve pre-anthesis traits adoption in Mediterranean bread wheat future breeding programs.

## Abbreviations

AGDM: above-ground dry matter
AN: anthesis
CTAB: cetyl trimethyl ammonium bromide
CW: chaff weight
DR: double ridge
DM: dry matter
EM: emergence
FL: flag leaf
FF: fertile floret
FFDM: fertile floret dry matter
FSR: floret survival rate (%)
GDD: growing degree days
GN: grain number
GPS: glume primordium stage
GW: grain weight
GY: grain yield
HD: heading
LS: leaf and spikelet initiation phase
MS: main stem
PH: plant height
RCBD: randomized complete block design
SDMa: spike dry matter at anthesis
SDMm: spike dry matter at maturity
SE: stem elongation phase
SF: sterile floret
TF: total florets
TL: tiller
TS: terminal spikelet
ZC: Zadoks’s code

## Acknowledgements

The authors are thankful to Israel Ministry of Agriculture and Rural Development for the ARO’s India-China 2-years Postdoctoral Fellowship to RR and masters’ scholarship to OZ. This research was supported by a grant from the Israeli Chief Scientist, Ministry of Agriculture (grant number: 20-10-00506) to DB, SA and RB-D. This research was also partly funded by the U.S. Agency for International Development of Middle East Research and Cooperation (grant number: SIS70017GR34037).

## Authorship contribution statement

**RR, OZ, GAS, DJB** and **RB-D** designed the scientific approach and experiments; **RR, DJB** and **OZ** performed the investigation, experiments, sampling and data collections; **KC** and **KN** assisted for sampling and data collection; **AC** assisted with genotypic screening; **RR** and **RB-D** analysed the data in software and worked on visualization; **RR, GAS** and **RB-D** wrote the manuscript; **KC, SA, DJB, GAS** and **RB-D** proof-read and edited the manuscript, and input improvement suggestions. All the authors approved the final version of the manuscript for submission.

## Supplementary Tables Caption

**Table S1.**
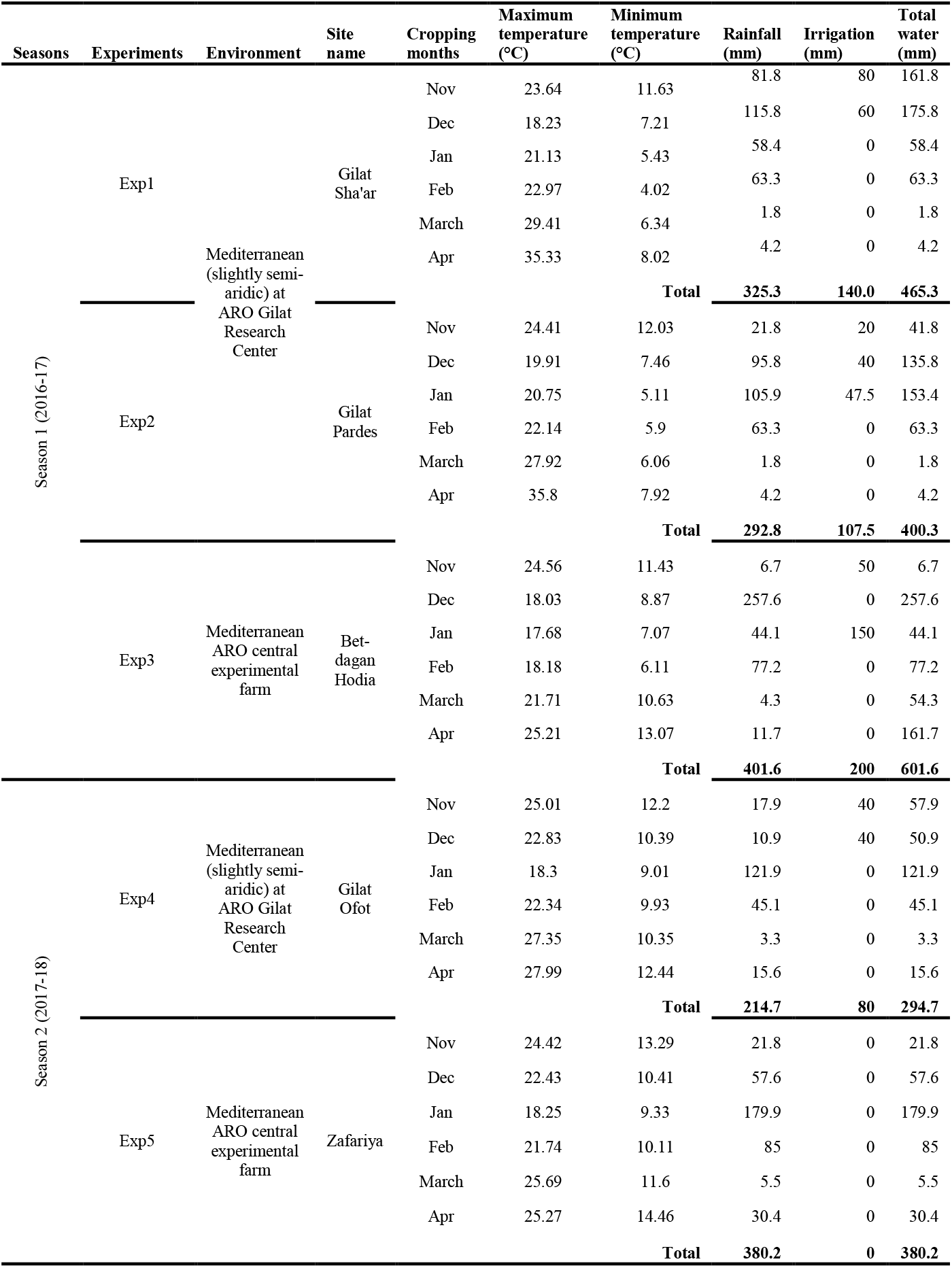
Temperature, rainfall and irrigation data for each cropping months across the five experimental sites

**Table S2.**
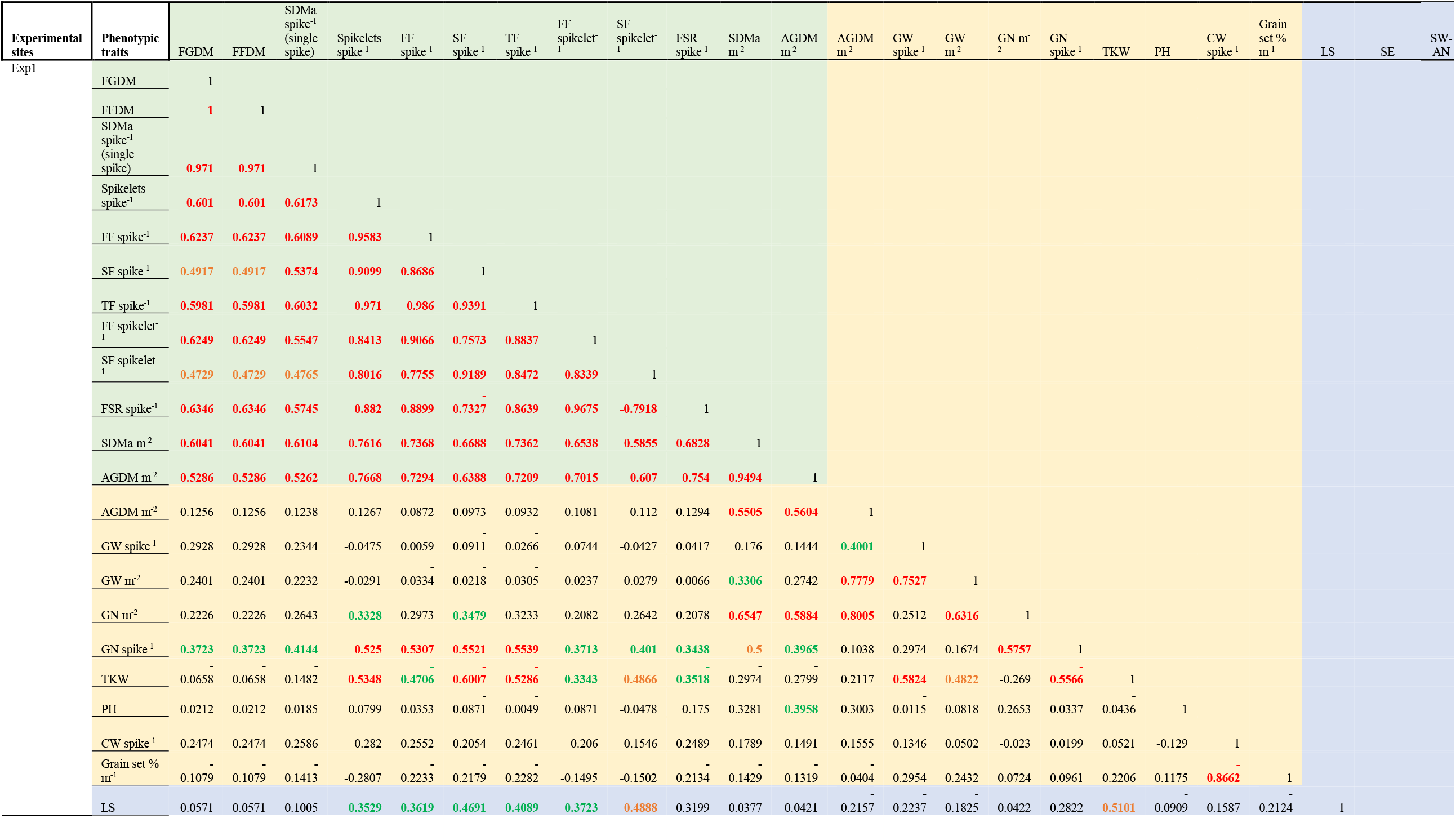
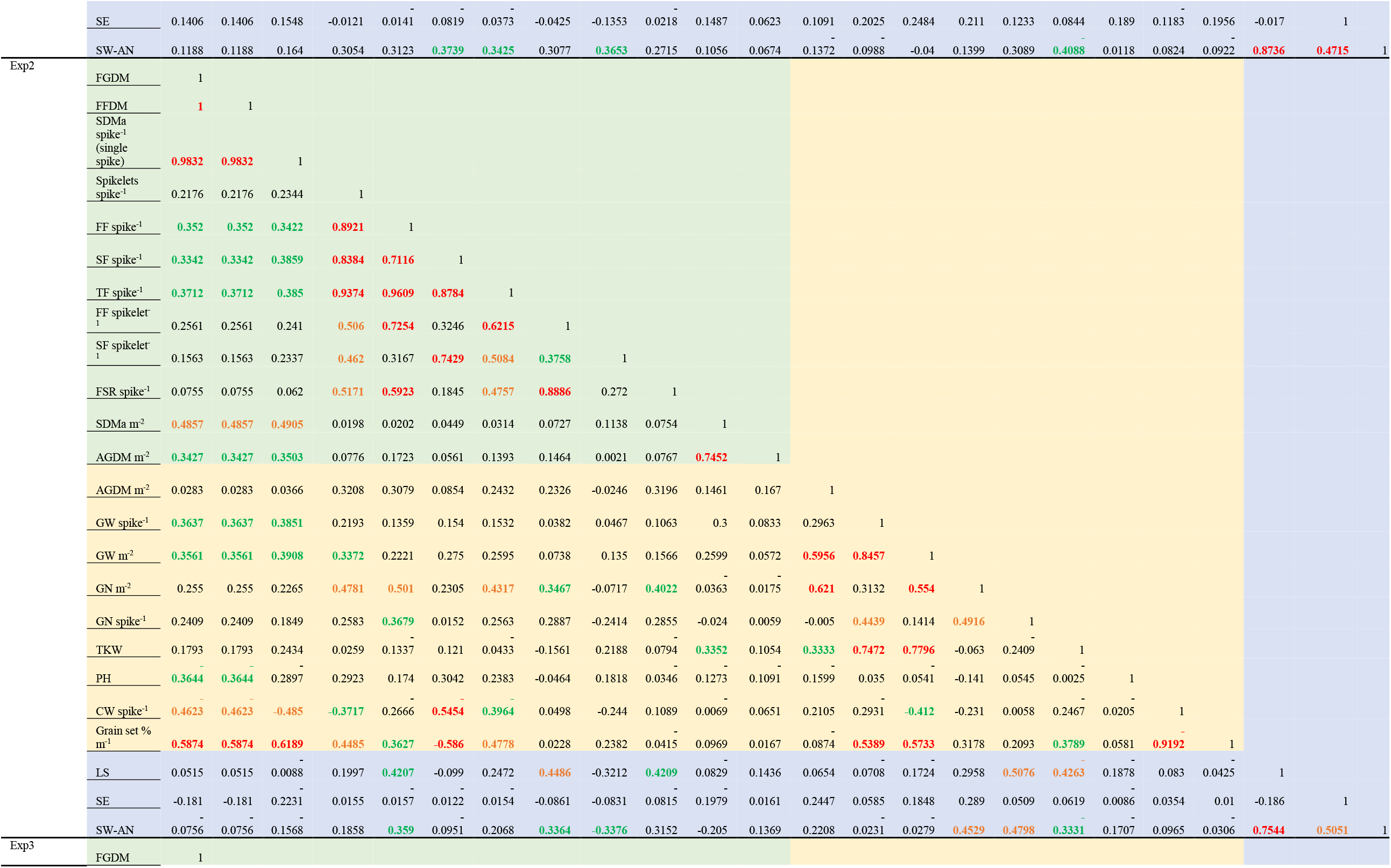
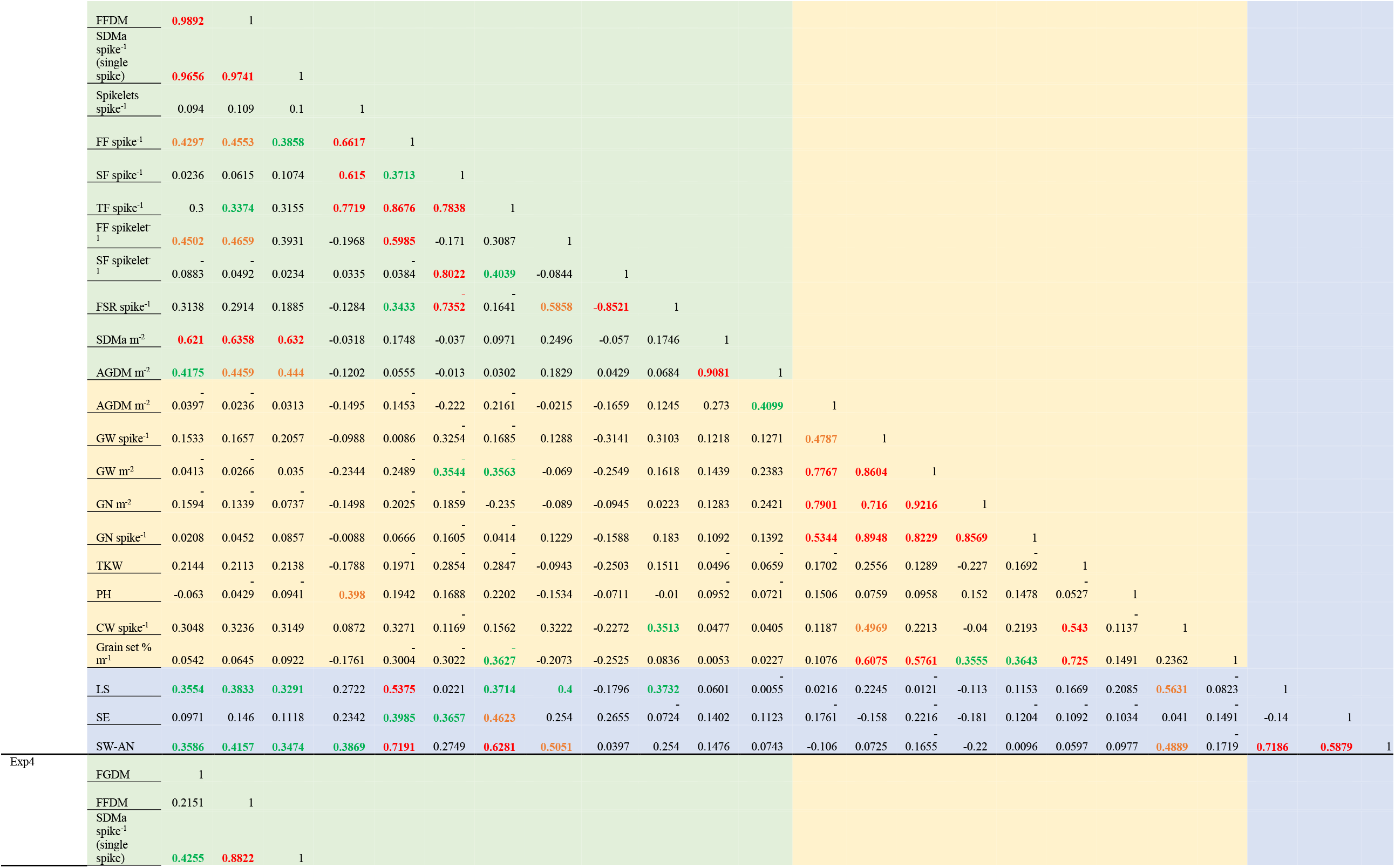
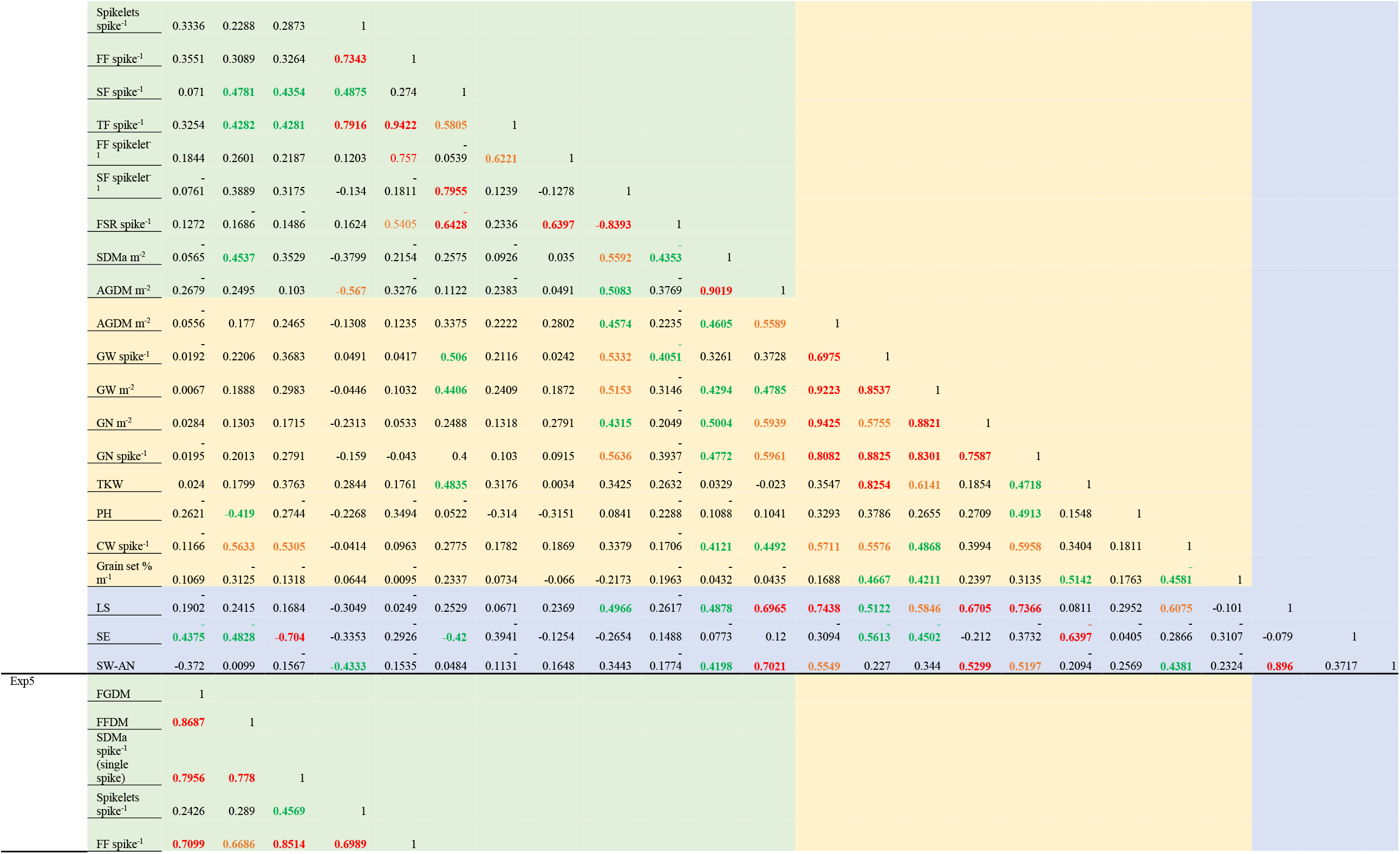
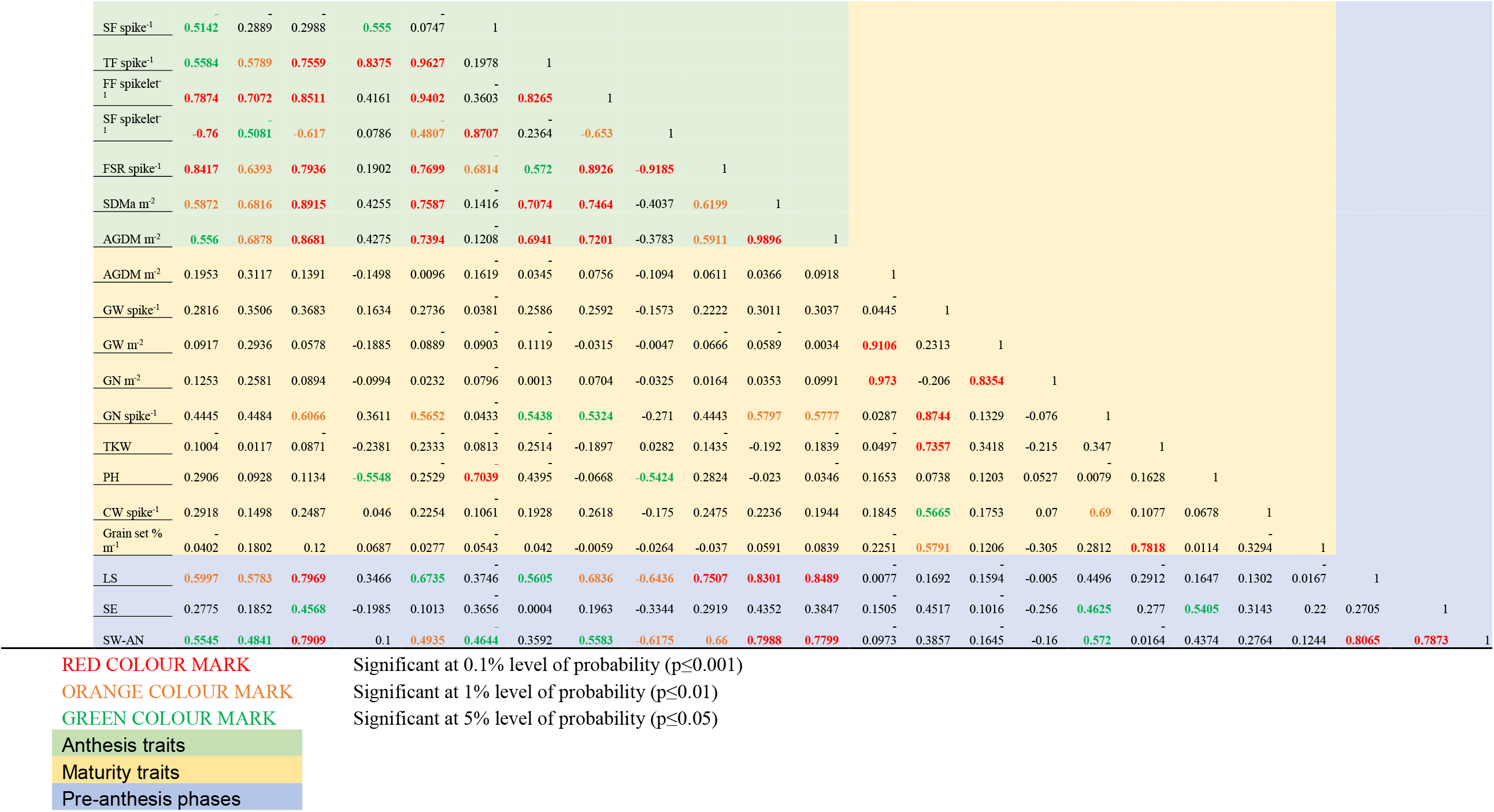
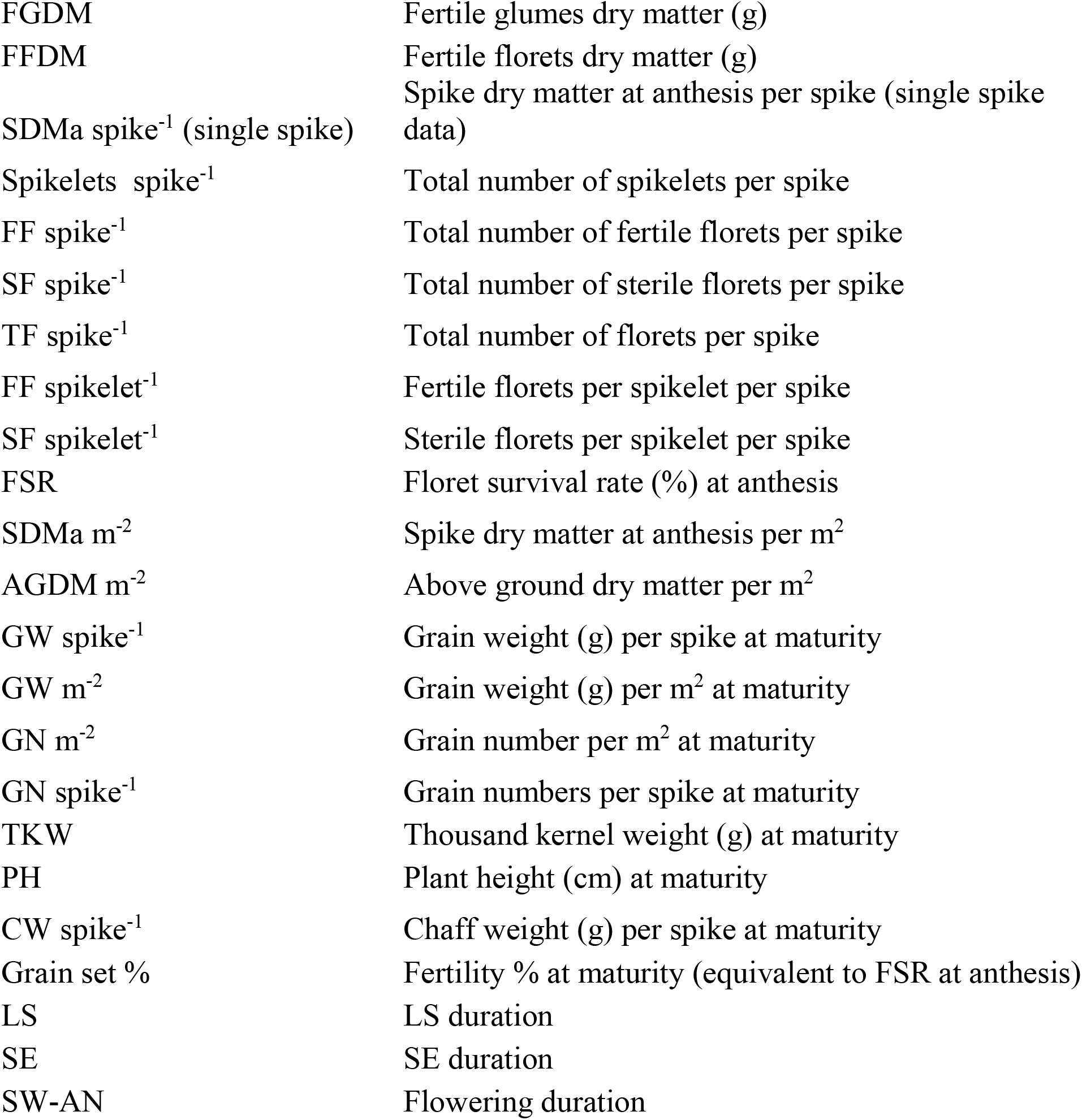
Correlation matrix for the phenotypic traits at both anthesis and maturity in five environments separately. Values of correlation coefficient (r) for each trait is represented in the matrix.

**Table S3.**
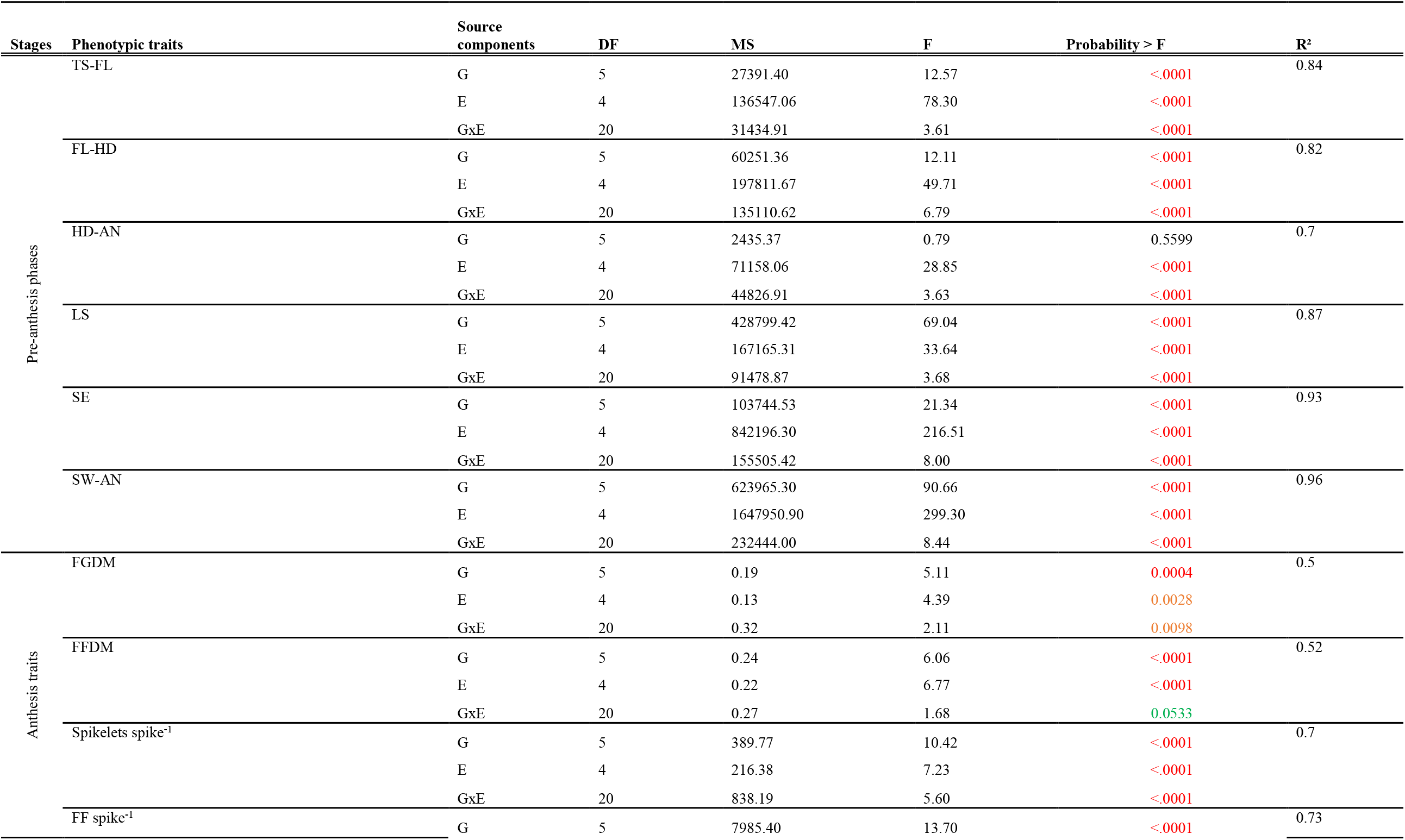
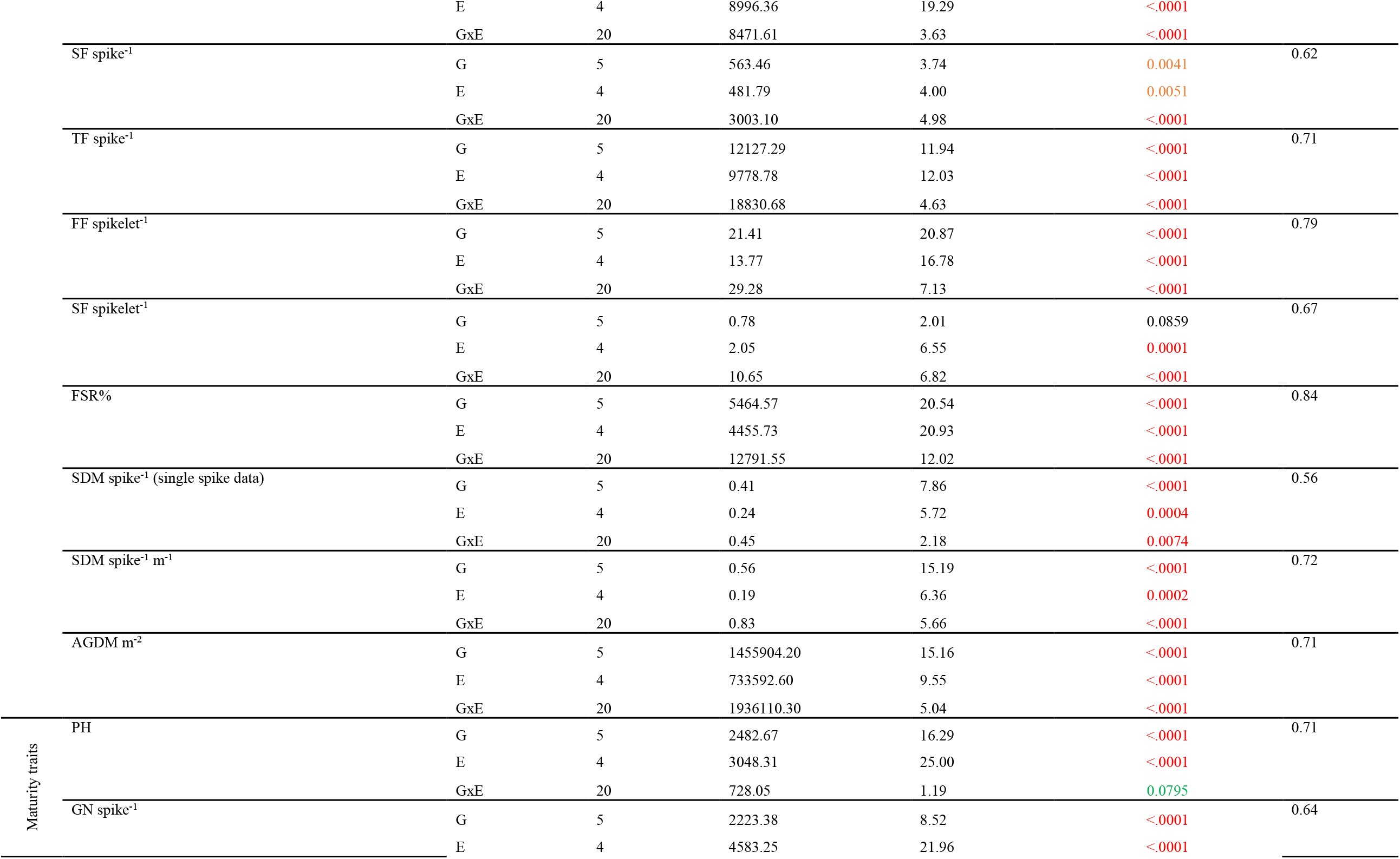
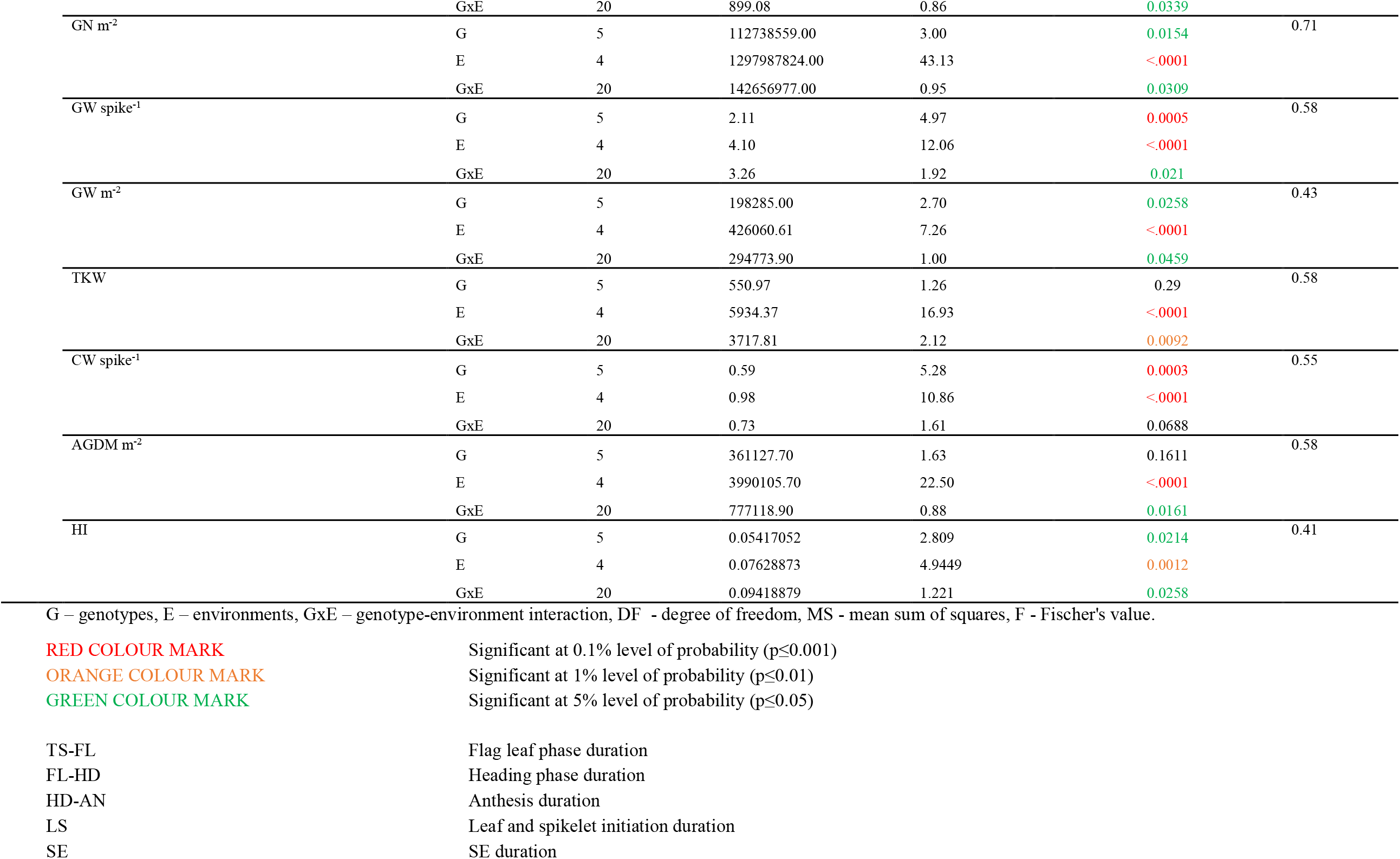
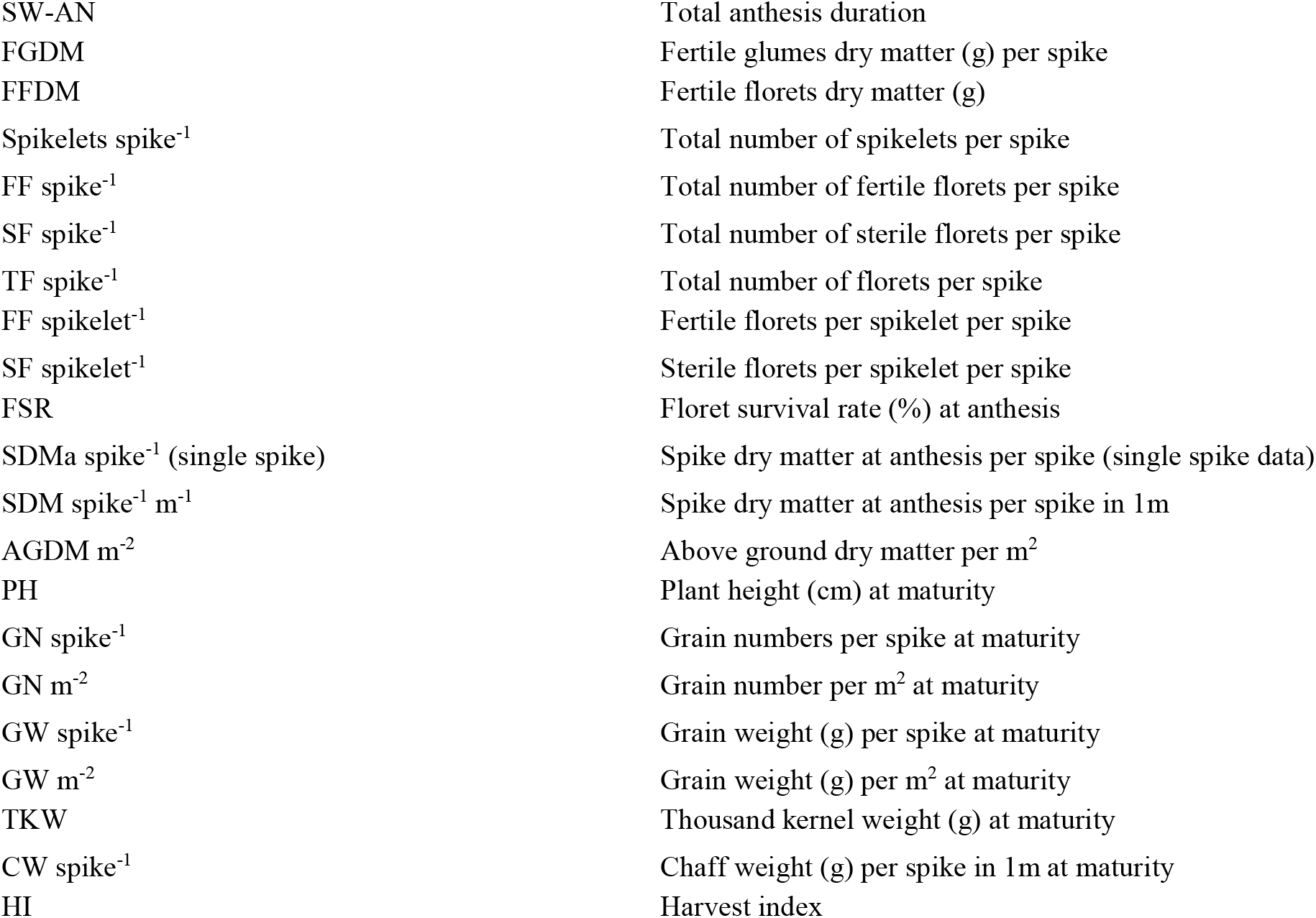
ANOVA for genotype-environment interaction (GxE) for pre-anthesis phases along with anthesis and maturity traits in 2016-17 and 2017-18.

## Supplementary Figures

**Figure S1.**
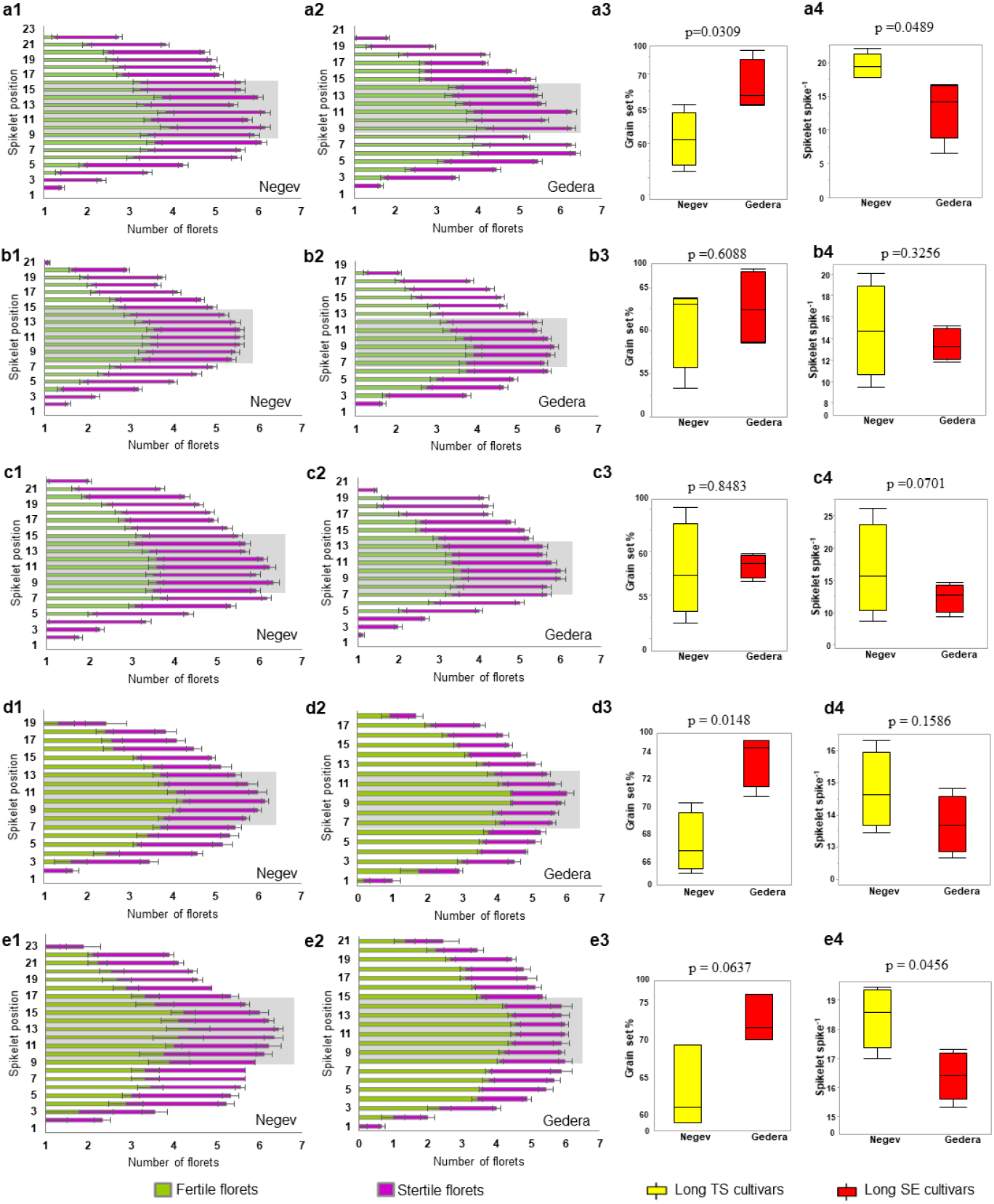
Floret conditions in the spike between the cultivar pair - Negev and Gedera. Number of grains (results of fertile florets at anthesis) and sterile florets of the cultivar pairs are representing as the bar graph according to each spikelet positions from the basal to apical part of the rachis node of each main-shoot spike at maturity. **a** represents Exp1, **b** represents Exp2, **c** represents Exp3, **d** represents Exp4, and **e** represents Exp5. Grey shaded part represents the position of central spikelets. The boxplots are representing the range of grain set % of the central spikelets at maturity (a3, b3, c3, d3, e3) and the mean spikelet number spike^-1^ including both main-shoot and tillers at maturity (a4, b4, c4, d4, e4) of Negev-Gedera pair in all five environments. The ‘*t*’ values for the significance probability between the pair is given in each plot.

**Figure S2.**
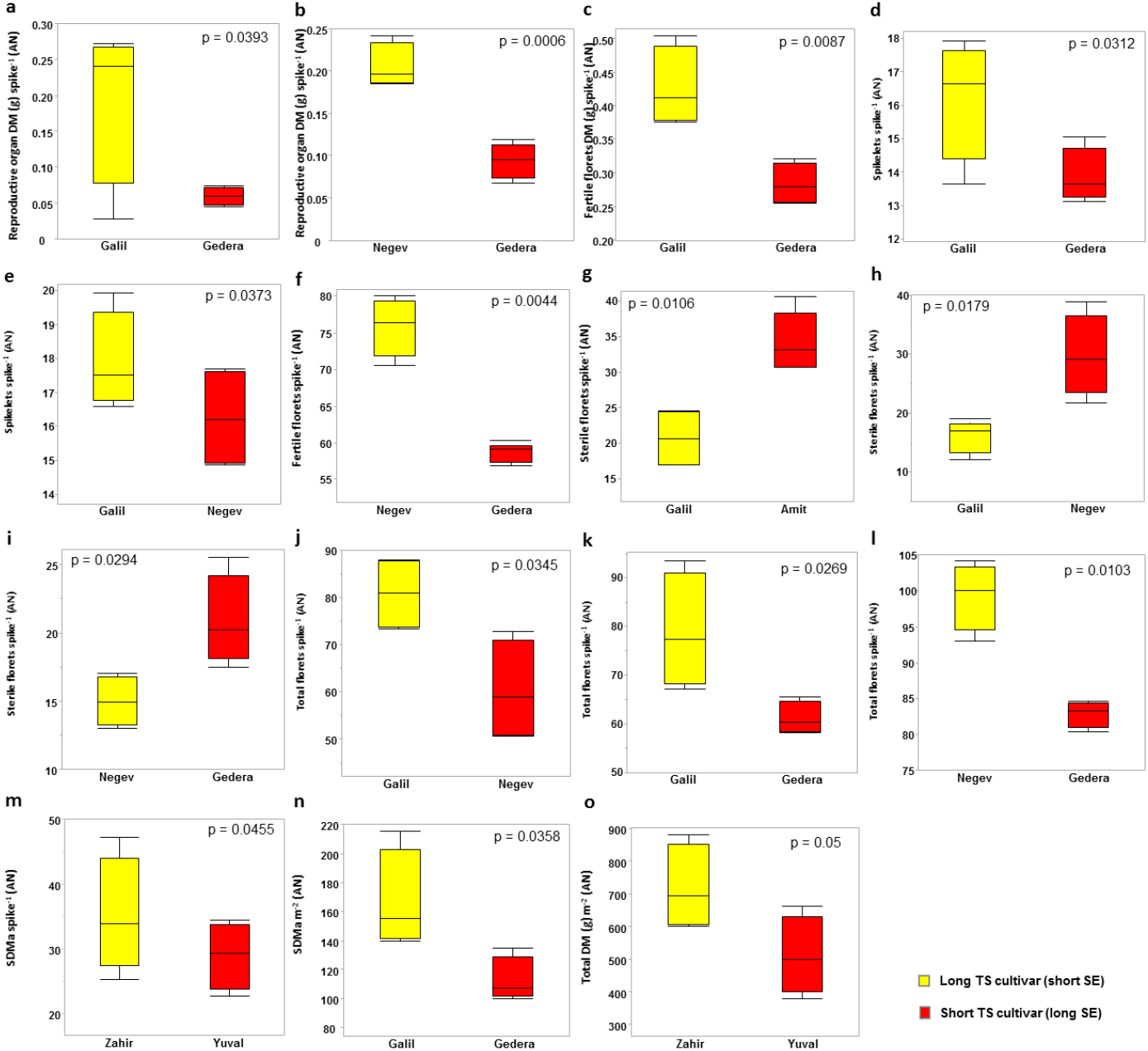
**Contrast analysis of spike traits and dry matter at anthesis** between cultivar pairs in a specific environment, which showed significant differences in LS with non-significant flowering time. Box plot representing reproductive organ dry matter spike^-1^ in Exp4 (a) and Exp5 (b), fertile florets dry matter spike^-1^ in Exp4 (c), spikelet numbers spike^-1^ in Exp4 (d) and Exp3 (e), fertile florets spike^-1^ in Exp5 (f), sterile florets spike^-1^ in Exp1 (g), Exp3 (h) and Exp4 (i), total florets spike^-1^ in Exp3 (j), Exp4 (k) and Exp5 (l), SDMa spike^-1^ in Exp3 (m), SDMa m^-2^ in Exp4 (n), above ground dry matter m^-2^ in Exp3 (o). The representing environments are Exp1 (Gilat Sha’ar), Exp3 (Bet-dagan Hodia), Exp4 (Gilat Ofot) and Exp5 (Zafariya). Representing all the spike traits and dry matter at anthesis are significant from 0.1% (p≤0.001) to 5% (p≤0.05) level of probability in the contrast analysis.

**Figure S3.**
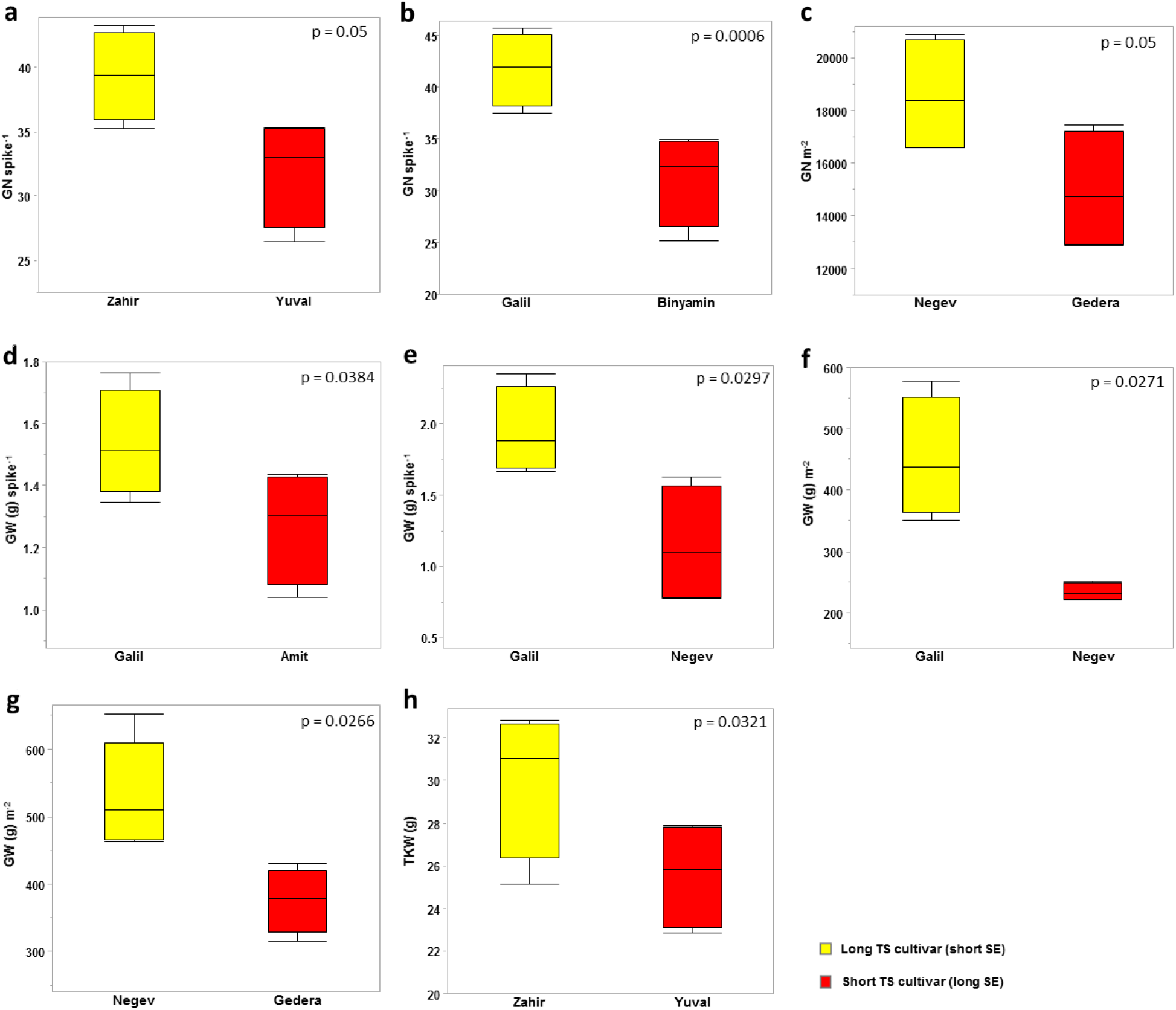
**Contrast analysis of yield components at maturity** between cultivar pairs in a specific environment, which showed significant differences in LS with non-significant flowering time. Box plot representing grain number spike^-1^ in Exp2 (a-b), grain number m^-2^ in Exp4 (c), grain weight spike^-1^ in Exp1 (d), Exp3 (e), grain weight m^-2^ in Exp3 (f) and Exp4 (g), thousand kernel weight (or TKW) in Exp4 (h). The representing environments are Exp1 (Gilat Sha’ar), Exp2 (Gilat Pardes), Exp3 (Bet-dagan Hodia) and Exp4 (Gilat Ofot). Representing all the spike traits and yield components at maturity are significant from 0.1% (p≤0.001) to 5% (p≤0.05) level of probability in the contrast analysis.

